# Quantitative spectral Linear Unmixing and Ratiometric FRET for live-cell imaging of protein interactions

**DOI:** 10.1101/2025.09.02.673709

**Authors:** Sonal Prasad

**Affiliations:** Cellular Neurophysiology, Center of Physiology, Hannover Medical School, 30625 Hannover, Germany; Department of Biomedical and Clinical Sciences, Linköping University, SE-581 83 Linköping, Sweden

**Keywords:** Lux-FRET, Ratiometric FRET, Spectral FRET, Quantitative confocal microscopy, cAMP biosensor, Serotonin receptors

## Abstract

We present a biophysical imaging strategy based on linear unmixing Förster resonance energy transfer (lux-FRET) for investigating protein-protein interactions and receptor-mediated signaling in live cells. This method utilizes spectral unmixing of FRET signals acquired via confocal laser scanning microscopy (LSM), enabling high-resolution quantification of molecular interactions with both spatial and temporal precision. Applying lux-FRET, we examined receptor-receptor interactions and downstream signaling events, including agonist specificity for 5-HT receptors. Ratiometric FRET measurements with a genetically encoded cAMP biosensor allowed us to assess biosensor sensitivity to cyclic nucleotides and receptor efficacy. Additionally, we explored physiological interactions between CD44 and 5-HT receptors and characterized the oligomerization state of the 5-HT1A receptor through apparent FRET efficiency analysis. Our findings demonstrate the utility of lux-FRET combined with quantitative molecular microscopy as a powerful tool for dissecting dynamic signaling mechanisms in live cells. This approach offers broad applicability for researchers studying receptor pharmacology, cellular signaling, and protein interaction dynamics.

**RESEARCH HIGHLIGHT:** We present a real-time imaging strategy combining lux-FRET with quantitative molecular microscopy to study protein interactions and receptor signaling in living cells. Using spectral and ratiometric FRET analysis, this method enables high-resolution visualization of dynamic molecular processes under physiological conditions.

**GRAPHICAL ABSTRACT:** 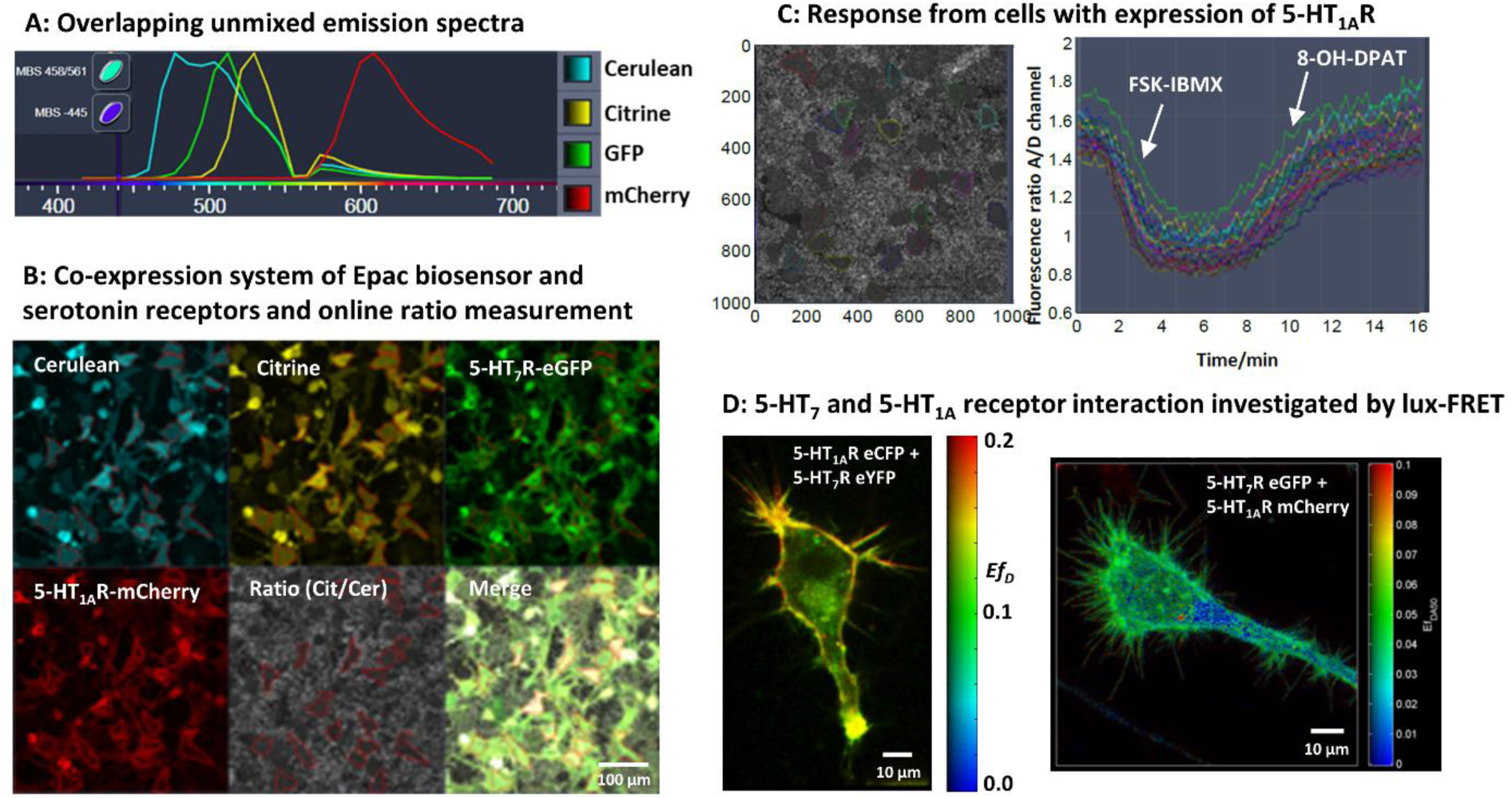

## INTRODUCTION

This study explores the application of linear unmixing FRET (lux-FRET) to investigate cAMP signaling and protein oligomerization in living cells, focusing on agonist specificity, 5-HT1A receptor mutants, and interactions between CD44 and serotonin receptors 5-HT1A and 5-HT7. Each result section highlights a distinct biological application of lux-FRET, demonstrating its utility in resolving receptor interactions and signaling dynamics at high spatial and temporal resolution.

Cyclic nucleotides such as cAMP, cGMP, cCMP, and cUMP are key second messengers that mediate cellular responses to hormones and neurotransmitters (Wolter, Golombek, & Seifert, 2011). Among these, cAMP and cGMP are well-characterized, synthesized by adenylyl and guanylyl cyclases (Beste & Seifert, 2013), and activate PKA and PKG, respectively. cAMP plays a central role in intracellular signal transduction, acting on effectors such as PKA, guanine exchange factors, PDEs, and cyclic nucleotide-gated ion channels (Taylor et al., 2004)(Francis, Busch, Corbin, & Sibley, 2010). To monitor cAMP dynamics in live cells, we used a FRET-based Epac biosensor, which enables consistent, real-time measurement under physiological conditions (Klarenbeek & Jalink, 2014; van der Krogt, Ogink, Ponsioen, & Jalink, 2008) (Gloerich & Bos, 2010).

Serotonin receptors 5-HT7R and 5-HT1AR modulate adenylyl cyclase activity via Gs and Gi proteins, respectively, resulting in opposing effects on cAMP levels—5-HT7R increases cAMP (Adham et al., 1998; Wirth et al., 2017), while 5-HT1AR decreases it (Albert et al., 1999; Kvachnina, 2005). These receptors exhibit distinct agonist affinities, with 5-HT7R responding strongly to 5-CT and moderately to 8-OH-DPAT. Selective targeting of 5-HT7R has shown therapeutic potential in mood and cognitive disorders (Speranza et al., 2017) (Hedlund et al., 2010). In co-expression experiments, we observed heterogeneous cAMP responses that could not be explained by receptor concentration alone. To investigate this, we applied a lux-FRET-based strategy to assess the impact of GPCR oligomerization on downstream signaling (Prasad et al., 2019).

We also examined the role of PKCα, encoded by the *PRKCA* gene, a membrane-associated kinase that regulates cellular functions through phosphorylation. Its activity is modulated by membrane interactions (Sun, Hu, Moradel, Weight, & Zhang, 2003) (Liu et al., 2017) and can be pharmacologically activated by phorbol 12-myristate 13-acetate (PMA), a potent tumor promoter that mimics the natural PKC activator diacylglycerol (Davis et al., 2018) (Mogha, Guariglia, Debata, Wen, & Banerjee, 2012).

Additionally, we explored the function of CD44, a transmembrane glycoprotein involved in cell adhesion, migration, and neuronal development through interactions with hyaluronic acid and matrix metalloproteinases (Senbanjo & Chellaiah, 2017). CD44 is critical for dendritic arborization and spine morphology (Bijata et al., 2017), and its inhibition produces neuronal changes similar to those induced by 5-HT7R activation, suggesting a functional link between CD44 and serotonergic signaling(Roszkowska et al., 2016).

Finally, we investigated 5-HT1A receptor oligomerization, a known feature of class A GPCRs, which can form homomers and heteromers (Renner et al., 2012). While oligomerization is well-documented, its physiological role remains incompletely understood. The 5-HT1AR has shown variable oligomerization behavior (Kobe et al., 2008; Renner et al., 2012; Woehler, Wlodarczyk, & Ponimaskin, 2009), and site-directed mutagenesis targeting intracellular regions has provided insights into the structural basis of receptor dimerization (Gorinski et al., 2012).

## METHODS

### Fluorescence spectroscopy

#### Spectral lux-FRET analysis in living cells

Spectroscopic measurements were performed using fluorescence spectrometer Fluorolog-322 (Horiba Jobin Yvon, Munich, Germany). This system is highly suitable for FRET investigations in lysed cells and cells in suspension for titration and time lapse experiments.

For quantitative measurements of cyclic nucleotides, titration experiments were performed in mouse N1E-115 neuroblastoma (N1E) cells transfected with plasmid DNA encoding 1 µg of CEpac biosensor. The cAMP-sensing Epac is a biosensor based on a FRET pair of the fluorophores eCFP and eYFP (Ponsioen et al., 2004). 16 h after transfection cells were washed and lysed in buffer (150mM NaCl, 10 mM HEPES and 1% Triton at pH 7.4) and washed three times with PBS and resuspended in buffer (150 mM NaCl, 5 mM KCl, 1 mM MgCl_2_.6H_2_O, 2 mM CaCl_2_.2H_2_O and 10 mM HEPES at pH 7.4 and 342 mOsmol) and homogenized and centrifuged. To inhibit the enzymatic activity of phosphodiesterases, 50 µM of phosphodiesterase inhibitor IBMX was added to the lysate. The cyclic nucleotides were titrated into the supernatant solution of the protein lysate. 2 ml of the supernatant was directly filled into quartz cuvettes equipped with a magnetic stirrer. Measurements were performed in 10 mm pathway quartz cuvettes at 1 nm wavelength steps with 4 nm spectral resolution for excitation and 2 nm for emission and integration time 1 s, continuously mixed by a magnetic stirrer. To suppress scattering and reabsorption, spectra were measured in a front face arrangement. To estimate the spectral contributions of light scattering and autofluorescence of cells, reference spectra of empty vector transfected cells were recorded and considered an additional background component for the fitting procedure. The chamber temperature was set to 25°C. For reference measurements, cells were co-transfected with plasmids encoding a single fluorophores eCFP and eYFP (only donor and only acceptor). To maintain total plasmid concentration, an equal amount of empty vector was transfected together with receptor. To estimate the spectral contributions autofluorescence of cells, reference spectra of empty vector transfected cells were used. The fluorescence excitation and emission spectra of eCFP and eYFP were obtained separately. The reference emission spectra of eCFP and eYFP used for the unmixing procedure were obtained at 440 nm excitation where the relative excitation of eCFP to eYFP was optimal. Background and Raman spectra were unmixed along with references during the data analysis. A CFP–YFP tandem construct with a 1:1 stoichiometry and E = 35 ± 3% was used to obtain R_TC_ and as positive control. As negative control, we used eCFP and eYFP, which are expressed in cytosol. Detailed information of FRET data evaluation is described (Prasad et., 2013) (Salonikidis, Zeug, Kobe, Ponimaskin, & Richter, 2008).

To find best excitation wavelengths for eCFP–eYFP pairs, we tested linear unmixing (lux-FRET) accuracy and error propagation by using various pairs of excitation wavelengths (440 nm, 458 nm, 488 nm). The recordings were done at three different excitations: 440 nm, 458 nm and 488 nm. We chose two wavelengths alternately for excitation. The emission was recorded at 450-600 nm for exc1: 440 nm, 460-600 nm for exc2: 458 nm and 480-600 nm for exc3: 488 nm. The acquisition protocol was kept same as described (Salonikidis et al., 2011; Wlodarczyk et al., 2008). For a comprehensive analysis of the emission properties of CEpac, its fluorescence spectrum was found as a linear combination of the fluorescence spectra of eCFP and eYFP in the complete concentration range of cAMP (0-1 mM) and for pH range (pH 6.5 – 8). We used spectral based alternate two excitation wavelength lux-FRET method to obtain a reliable measure of the absolute cAMP concentrations as previously described (Wlodarczyk et al., 2008).

We chose N1E cells as a model since they are known not to express any G-protein coupled 5-HT receptors. For eGFP-mCherry FRET pairs, cells were co-transfected at 1:1 ratio with plasmid CD44 and 5-HT receptors tagged with eGFP and mCherry fluorophores. The excitation wavelengths for donor 480 nm and acceptor 580 nm was selected after screening from six excitations exc1 (470 nm-490 nm) and exc2 (570 nm-590 nm) which yielded higher apparent FRET efficiency.

For the time-course experiments cells were co-transfected with 5-HT1AR WT and/or 5-HT1AR mutatant plasmids tagged with eCFP and eYFP fluorophores. After 16h, cells were resuspended in PBS and spectroscopic measurements. During the experiment, two emission spectra were obtained for each time point at excitations 458 nm and 488 nm with 5 nm of spectral resolution for excitation and emission, respectively. Cells expressing only single fluorophores were used for reference. The acquisition and experimental protocol were followed as described (Gorinski et al., 2012). We determined the apparent FRET efficiencies *Ef_D_* and *Ef_A_* for WT and mutants of the 5-HT1AR, by using lux-FRET method described (Wlodarczyk et al., 2008). Quantitative comparison of FRET values was obtained at different donor to acceptor ratios by keeping the total concentration of plasmids encoding for donor and acceptor constant. We fitted the model characterizing apparent FRET efficiency (*Ef_D_*), as a function of donor mole fraction (*x_D_*), for oligomeric structures (Veatch & Stryer, 1977) with a slight addition for use with *Ef_A_* according to (Meyer et al., 2006) to our experimental data. The true transfer efficiency, E, and the number of units, n, interacting in the oligomeric complex was estimated from the dimerization model fitting.

### Fluorescence microscopy

#### Time series measurement for cAMP responses

All data were acquired on a Zeiss LSM 780 controlled with ZEN 2012 software. Confocal imaging was performed after comprehensive calibration of the system according to (Butzlaff et al., 2015; Prasad et al., 2013). The following acquisition configuration was used: 40x/1.2 NA water immersion objective, bit depth 16-bit, and pinhole of 1.7 AU at zoom 0.6. N1E cells were subjected to fluorescence ratio time series measurements to monitor cAMP responses with an optimized variant of FRET-based Epac biosensor (Salonikidis et al., 2011). This approach provides information about relative changes in cAMP concentrations with good spatial and temporal resolution. A conformational change of the cAMP bisensor CEpac-construct (Cerulean-Epac(δDEP-CD)-Citrine) is induced by cAMP binding to the regulatory domain of exchange proteins activated by cAMP (Epac). The distance between the two fluorophores increases after cAMP-binding, resulting in decrease in FRET intensity. CEpac acts in an inverse manner, showing high FRET at low cAMP concentration and low FRET at high cAMP concentration. Decrease in cAMP leads to an increase in the energy transfer between CFP variant Cerulean (D) and YFP variant Citrine (A), whereas increase in cAMP diminishes the energy transfer. The measurement of the donor/acceptor fluorescence emission ratio (A/D ratio) in the tandem biosensor varies with the changes in cAMP concentration when excited at the donor excitation wavelength. The standard ratiometric readout for FRET-based biosensors, the A/D ratio does thus show a decrease in the A/D ratio upon increasing [cAMP]. Unless the A/D ratio shows apparently smaller amplitudes of ratio changes over the inverse, the donor/acceptor ratio, the A/D ratio was used to avoid division by small numbers at low FRET.

Cells expressing CEpac biosensor and 5-HT7R-eGFP were excited with 440 nm and 5 HT1AR-mCherry with 561 nm laser line (MBS 440 and MBS 458/561, emission range 450 – 600 nm). Images were acquired at a frame size of 1024×1024 pixels and a frame acquisition time of 6.8 s. During the experiments, 100 cycles with 10 s intervals along with continuous refocusing using the Zeiss ‘Definite Focus’ were used. The fluorescence spectra of Cerulean, Citrine, eGFP and mCherry from single transfections, acquired in lambda mode and background corrected, were used as references for unmixing in the online fingerprinting mode (Fig. S9A). Cells that showed specific plasma membrane labeled with moderate 5-HT7R and 5-HT1AR and moderate cAMP biosensor expression were selected for analysis. In the time series experiments, 18–20 frames (3.0 min) were captured prior to application. Agonists and antagonists were applied via a perfusion system with a constant flow rate of 1.0 ml min−1, thus enabling a complete exchange of solution within 1.0 min. The application protocol was adjusted by optimizing the agonist and antagonist concentrations and the blocking time. Working concentrations were 20 µM for LP-211 and 5-HT, 100 nM for SB, 10 nM for WAY, 5 µM for Forskolin (FSK), and 50 µM for IBMX. Concentrations for FSK and IBMX were chosen to obtain a cAMP increase high enough to show significant cAMP decrease by activating 5-HT1AR, but low enough not to saturate the biosensor and not suppress the 5-HT1AR signaling. The blocking time was set to 10 min prior to agonist application. Ratiometric quantitative FRET data evaluation was conducted offline (Fig. S9B) (Prasad et al., 2019). The intensities in all four colors were determined from whole-cell regions of interest (ROIs) drawn manually at a higher magnification online in parallel to ongoing measurements. The measurement of the acceptor/ donor emission ratio in the tandem biosensor (1:1 donor/acceptor stoichiometry) was excited at the donor excitation wavelength.

Experiments were designed accordingly to rule out any kind of quenching effects from (transient) over-expression of receptor pairs simply soaking up Gα/Gßγ subunits which need to be considered. We found no correlation whatsoever between the receptor intensity and cAMP response for the concentration range used in this study.

Fitting models for A/D ratio time series data: Since 5-HT7R and 5-HT1AR are known to regulate the intracellular cAMP production upon activation, mathematical modeling was used to quantify the change in intracellular cAMP. Three fit models were developed to describe the time-dependent changes in the cAMP concentration: fit model no. 1 – a single exponential fit with a polynomial offset (*polyoff*), which delivers decay time *τ1* and amplitude *A1* starting at time point *t0*; fit model no. 2 – a two exponential fit with polynomial offset at two time points (*t1* and *t2*), which delivers a set of two amplitude (*A1* and *A2*) and decay time (*τ1* and*τ2*) parameters for cAMP up regulation and down regulation; and fit model no. 3 – a double exponential fit, which combines two biological processes at one time point. Model no. 3 also delivers two sets of parameters of amplitude and decay time values, where the real value of the decay time and amplitude cannot be deduced due to the complexities of model fitting. Thus, raw data for the third type of response were simply fitted with the model, and the deduced quantities could rarely be used. The resulting amplitudes from the exponential fit models must be read as cAMP increase for *A1*<0 and a cAMP decrease for *A2*>0 as described (Prasad et al., 2019).

For cyclic nucleotides time series measurements, confocal imaging was performed on cells expressing 1 µg of CEpac biosensor excited with a 440 nm laser line (MBS 440, emission range 450-600 nm). Cells were pre-stimulated with 50 μM of IBMX. After 5 min of acquisition, within 15-20 s on application of 100 µM cNMP-AM’s to the bath, the change in the fluorescence intensity ratio of cAMP response was obtained in online fingerprinting mode and further processed offline.

In cells co-expressing cAMP biosensor and 5-HT1AR WT/mutants time series measurements were performed. Cells were stimulated with 5 µM FSK in combination with 50 µM IBMX and 20 µM 5-HT. To see significant cAMP decrease by activating 5-HT1AR signaling, high enough concentrations of FSK and IMBX were used. Ratiometric FRET data evaluation was conducted offline.

#### Lux-FRET measurement

Lux-FRET imaging was performed on cells that co-expressed eGFP/mCherry-tagged CD44 and eGFP/mCherry-tagged 5-HT7R or 5-HT1AR at a plasmid concentration of 1:1 ratio. Z-stacks (10 slices, 1.0 µm spacing) were acquired at 488 nm and 561 nm excitation, MBS 488/561, and an emission range 500-700 nm. All other acquisition settings and parameters were kept same as previously described (Prasad et al., 2019). We performed detailed pixel-based analysis and achieved a high spatial resolution in 3D in detail according to the protocol described (Butzlaff, Weigel, Ponimaskin, & Zeug, 2015; Prasad, Zeug, & Ponimaskin, 2013). Alternating line wise switching excitation was achieved with a custom-tailored acquisition protocol generated by the LIC Macro toolbox. The laser power was set to 3% for 488 nm and 561 nm excitations to avoid bleaching. The correlation between FRET parameters and intensity was used to verify that receptor concentration-related artifacts, such as ‘molecular crowding’, are avoided. For lux-FRET analysis, properties were obtained from all the regions of interest (ROIs). Since lux-FRET does also provide relative concentrations of donor and acceptor, it is possible to distinguish between the influence of interaction and concentration artifacts. Images for reference were acquired from cells that expressed fluorophores separately. The eGFP-mCherry tandem construct was used in the experiment with the previously described acquisition settings for a single image. We obtained *Ef_D_* and *𝐸f_A_* over 𝑥_𝐷_ (Bijata et al., 2017; Prasad et al., 2019), fitted with the dimerization model using the lux-FRET method (Renner et al., 2012; Wlodarczyk et al., 2008). During the time-course experiments, the cells were perfused with buffer for 5 min followed by 10 μM 5-CT for 15 min. For the lux-FRET study, we took special care to provide accurate data quality. For that, substantial amount of calibration work and controls were performed. In case of lux-FRET measurements, we performed controls with non-fretting membrane proteins (e.g. 5-HT7R and CD68) acting as negative control (data not shown).

For cells co-expressing eCFP-tagged PKCα and eYFP-tagged PKCα (0.5 µg each) serving as donor and acceptor molecules keeping 1 µg total concentration, we performed FRET measurement using two-track channel mode as previously described. Z-stacks with 10 frames were acquired at excitations 440 nm −lp 10% and 514 nm −lp 5%, MBS 440/514, zoom 0.1, pinhole 1.0 AU, and an emission range 450-600 nm before and after 10 minutes of application of PMA. 10 μM PMA was applied locally after 3 minutes from the start of the acquisition. The drug was dissolved in DMSO at 1 mM concentration with further dilution in buffer solution. Reference images were acquired from cells that separately expressed fluorophores and CFP-YFP tandem construct. Acquisition setting and parameters were kept same as previously described (Prasad et al., 2013).

### Data evaluation

Imaging data were evaluated offline using Matlab (Mathworks, Natick, MA, USA, licensed LiU). To determine the FRET efficiency for CD44.– 5-HTR interactions, we performed pixel-based lux-FRET analysis as described in detail previously (Prasad et al., 2013; Wlodarczyket al., 2008). Data where shift- and background-corrected and extracted from manually drawn ROIs covering the whole cell including the plasma membrane. The s.e.m was derived for all FRET quantities. The same ROIs were used for the lux-FRET analysis. Contour plots of the 2D histogram and maximum intensity Z-projections of unmixed images and apparent FRET efficiencies were generated in Matlab.

Three ratio models were applied to describe the time-depending changes in the cAMP concentration. The fluorescence ratio signal was normalized to 1.0 by dividing the ratios of each trace by its emission ratio immediately before stimulation (defined as ‘*offset*’). GraphPad Prism8.0 (licensed LiU) was used for illustration of fluorescence ratio traces.

## RESULTS

### Biophysical characterization of cNMPs using CEpac biosensor

We characterized the CEpac FRET-based biosensor in respect to its sensitivity for cyclic nucleotides (cAMP, cCMP, cGMP, cUMP) by performing titration measurements at concentration (0-1 mM) using spectrofluorometer. We found that cAMP activates the biosensor at 1 µM of concentration as seen from the decrease in fluorescence intensity of cAMP dependent emission spectrum and saturates at following higher concentrations (Fig. 1A). While the cyclic nucleotide cCMP caused low amplitude change of the CEpac biosensor at EC_50_ of 100 µM concentration (Fig. S1i) indicating a less specific binding to the Epac protein. Other cyclic nucleotides such as cGMP and cNMP showed no activation of the biosensor demonstrating weaker binding to the biosensor (Fig. S1ii, iii). The unmixing of the CEpac emission spectrum for cNMPs at 440 nm excitation fitted by a linear combination of the reference spectra for eCFP (D), eYFP (A), Raman, and background is shown (Fig. 1B; Fig. S2A). FRET efficiency (E) at four different combinations of the three excitation wavelengths was calculated for cNMPs and plotted as a function of concentration (Fig. 1C; Fig. S2B). The change in FRET efficiency of the biosensor caused by [cAMP] changes allowed quantitative measurement of cAMP concentration (Fig. 1D). We determined fluorescence ratio change (A/D) for all cNMP’s at 440 nm excitation which showed a sharp decrease in the fluorescence ratio at 1 µM for cAMP, and no change observed for other cNMP’s (Fig. 1E). To conclude, cAMP was the most effective activator of the biosensor.

**Figure 1.**
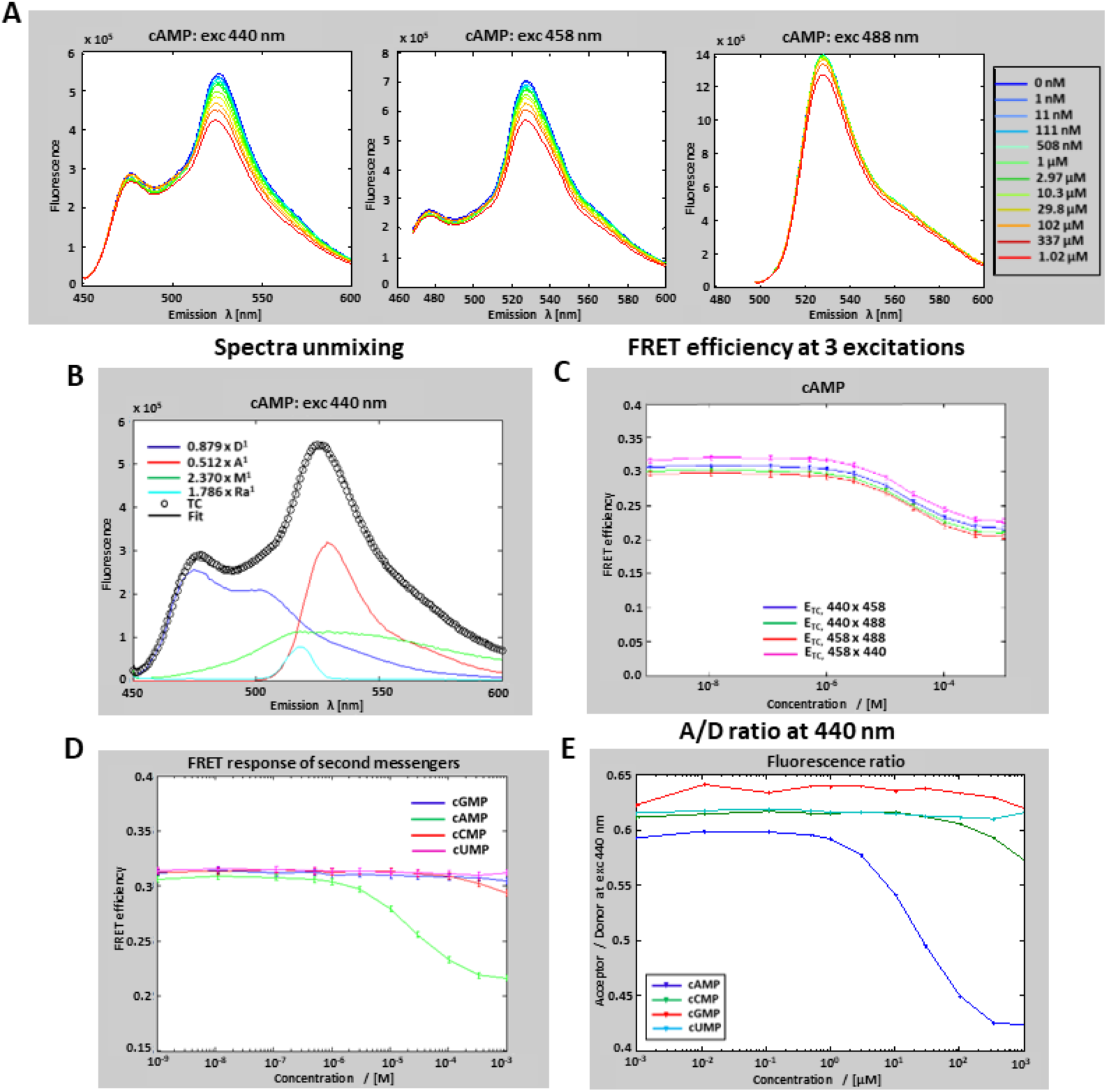
Cyclic nucleotides titration to cAMP biosensor investigated by lux-FRET. (A) [cAMP]-dependent Epac emission spectra were obtained from a diluted supernatant of homogenized and centrifuged cells transfected with Epac. The emission spectrum was obtained at 440 nm, 458 nm, 488 nm excitations showing change in the fluorescence emission signal at 1 µM. cAMP was directly applied into the supernatant solution. The gray arrow indicates the intensity change with increasing [cAMP]. (B) Epac emission spectra measured at 440 nm excitation was fitted by a linear combination of reference spectra for YFP, CFP, Raman, and background. The reference spectra were obtained separately. All presented spectra are corrected for background and autofluorescence. (C) FRET efficiency of cAMP at four different combinations of the three excitations. (D) Bar plot of FRET efficiency and fluorescence ratio (D/A) change (E) for second messengers at 440 nm excitation over a concentration range of 0-1 mM. Experiments were performed in lysed cells transiently expressing cAMP biosensor CEpac and repeated at least three times. See Figures S1 and S2 for other cyclic nucleotides.

Next, we performed time series measurements for cyclic nucleotide responses in N1E cells expressing CEpac biosensor. Highly membrane-permeant precursors of the second messenger cyclic nucleotides (cNMPs-AM) were applied locally to the cells. The cells were bathed in buffer solution consisting of IBMX, a non-selective cAMP and cGMP PDE inhibitor to block all the PDEs. The cNMP-AM’s esters are metabolically activable pro drug of the cNMP’s (Werner, Schwede, Genieser, Geiger, & Butt, 2011). The polar cAMP which is trapped inside the cell is released after permeation of the precursor and metabolic activation by esterases, subsequently metabolized quickly, resulting in a change in the fluorescence intensity ratio. Experiments were carried out with continuous perfusion of IBMX and manual application the cNMP-AM esters. The manual application of nucleotide esters could cause a slight perturbation, which is visible over the time-course fluorescence ratio data. These fluctuations show a drop and deep rise behaviour (artefact), which are clearly distinct from the stimulation response, see red vertical line in Fig. 2A.

**Figure 2.**
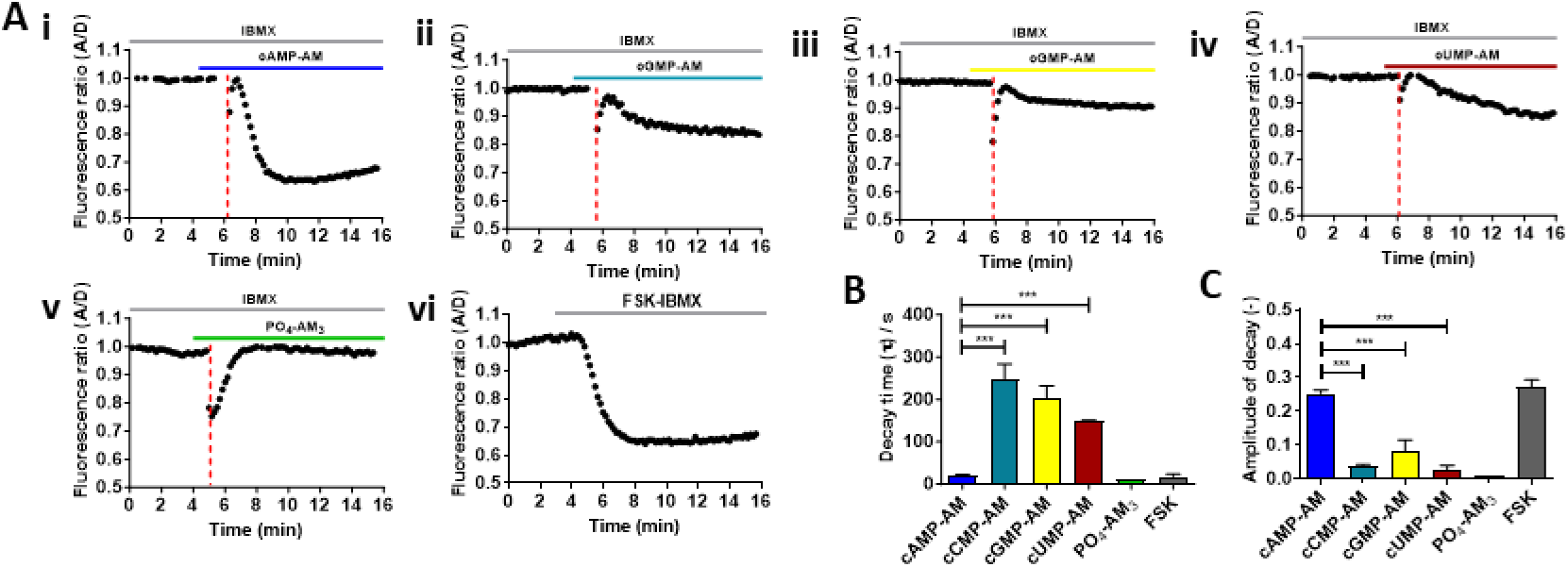
cNMP-AM’s induced changes in the fluorescence ratio correspond to cAMP biosensor in N1E cells. (A) Representative traces showing change in the fluorescence ratio caused upon bath application of 100 µM of cNMP-AM esters (i-iv) and 33 µM of PO_4_-AM_4_ control reagent (v) and FSK as a positive control (vi) in combination with continuous inhibition of phosphodiesterase with 50 µM IBMX (n≥30). Experiments were performed in cells transiently expressing cAMP biosensor CEpac. Initial values of the fluorescence ratio were normalized to 1. (B, C) The decay time and amplitude of the cAMP response in the presence of cNMPs-AM and applying fit model no1 described in (Prasad et al., 2019). Data are the means ± s.e.m of at least two separate experiments. ***P<0.001; n.s., not significant; one -way ANOVA (Bonferroni’s multiple comparisons test).

The cAMP response was recorded at a single-cell level by monitoring the CEpac fluorescence intensity ratio of acceptor over donor (A/D ratio). In the presence of cAMP-AM the fluorescence intensity ratio decreased significantly due to increase in the intracellular cAMP, response visible right after the artefact (Fig. 2Ai). There was no significant decrease in the fluorescence ratio on application of other cNMP’s-AM (Fig. 2Aii, iii, iv). 33 µM of PO_4_-AM_3_ reagent was used as a negative control showing no decrease in the fluorescence ratio (Fig. 2Av). Forskolin (FSK) was used as a positive control showing a large decrease in the fluorescence ratio signal (Fig. 2Avi). We excluded the artefact when selecting traces for fitting the data. Statistical analysis by fitting time dependence of the fluorescence ratio data to a single exponential fit model no 1 (Prasad, Ponimaskin, & Zeug, 2019) revealed that cAMP-AM had significantly shortest activation time and the largest amplitude of the cAMP response (τ = 21 ± 1.3 s, A = - 0.25 ± 0.01, N= 2, n=26; where N is the number of experiments and n is the number of cells) compared with cCMP-AM (τ = 241 ± 39 s, A = - 0.03 ± 0.05, N= 2, n=19), cGMP-AM (τ = 202 ± 29 s, A = - 0.02 ± 0.01, N= 2, n=14) and cUMP-AM (τ = 149 ± 1.2 s, A = - 0.02 ± 0.01, N= 12, n=3) (Fig. 2B and C; Bonferroni’s multiple comparison test).

Thus, data from Fig. 1 and 2 confirmed that CEpac biosensor was highly selective to cAMP and is activated upon binding of cAMP and no other second messenger cyclic nucleotides.

### Biophysical characterization of ligand LP-211 for 5-HT receptor activation

Biophysical characterization of agonist LP-211 to determine its specificity towards 5-HT receptors by measuring cAMP signals were performed in living cells. Fluorescence intensity ratio experiments were performed in N1E cells co-expressing CEpac FRET-based cAMP biosensor with 5-HT7R tagged eGFP or with 5-HT1AR tagged mCherry. The spectrally highly overlapping signals have been unmixed using the Zeiss LSM online fingerprinting mode (see methods). No bystander effects were observed from the overexpression system as the relative expression of each of the receptors were controlled in respect to the CEpac biosensor for experiments at single cell level. Downstream signaling kinetics of both receptors 5-HT7 and 5-HT1A, first individually, and in combination were characterized. The cAMP response was recorded by monitoring the biosensor fluorescence intensity ratio. The 5-HT receptors were activated using agonist 10 µM LP-211. In cells that expressed 5-HT7R, LP-211 was able to increase the intracellular cAMP level via Gs signaling (Fig. 3A, orange trace). The direct activation of 5-HT7R was shown as negative change of the A/D ratio change of the CEpac biosensor upon increase of cAMP. When cells were pretreated with the specific 5-HT7R antagonist SB-269970 (Fig. 3A, SB, green trace), the cAMP response was fully blocked. Statistical analysis by fitting the time dependence of the A/D ratio data to a single exponential fit model no. 1 (Prasad et al., 2019) revealed short activation time and significantly large amplitude of the cAMP response (*τ1*=56 ± 3.0 s, *A1*=-0.28 ± 0.01, N=2, n=30) (Fig. 3B).

**Figure 3.**
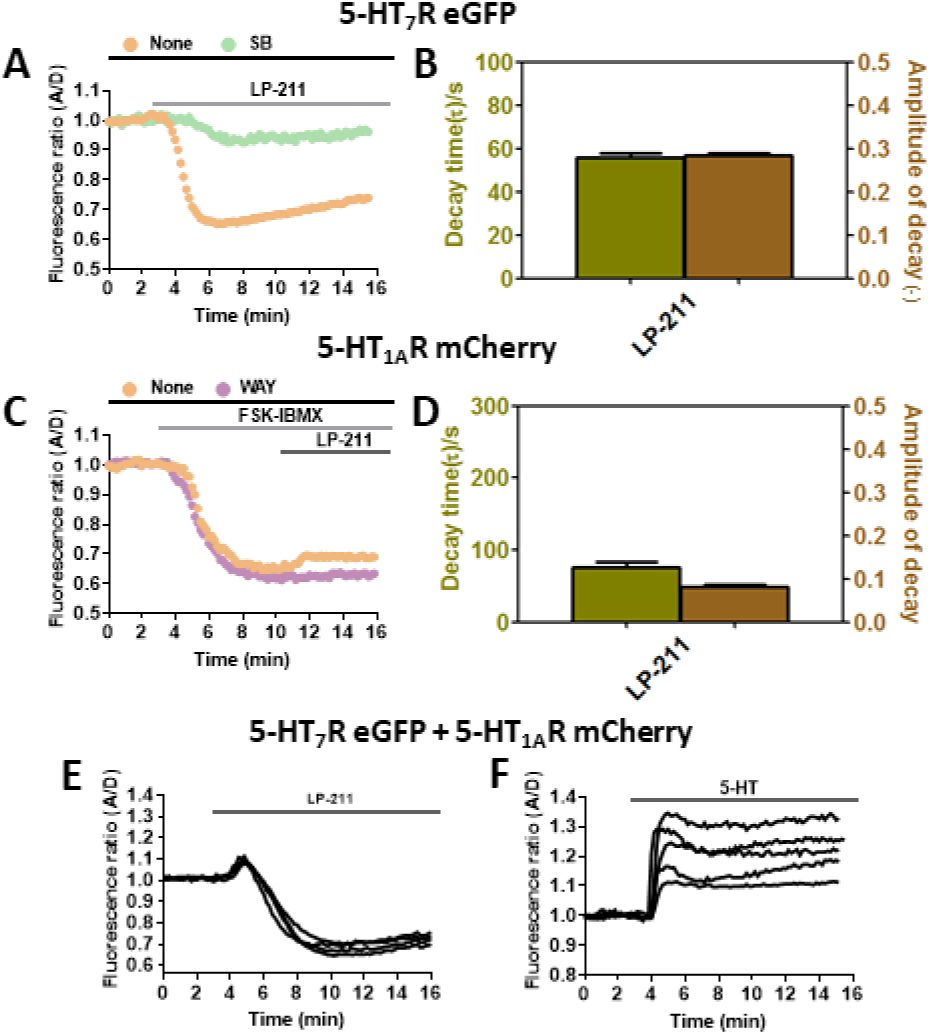
LP-211 induced change in the fluorescence ratio correspond to receptor activation in N1E cells. (A) Representative traces of the fluorescence ratio in response to agonist 10 µM LP-211 while pre-blocking 5-HT7R with SB (n≥30). (B) The decay time and amplitude of the cAMP response induced by 5-HT7R stimulation with LP-211 in absence of SB and applying fit model no1 described in (Prasad et al., 2019). (C) Representative traces of the fluorescence ratio in response to agonist LP-211 while pre-blocking 5-HT1AR with WAY in combination with FSK-IBMX stimulation (n≥30). (D) The decay time and amplitude of the cAMP response induced by 5-HT1AR stimulation with LP-211 in absence of WAY and applying fit model no2 described in (Prasad et al., 2019). Experiments were performed in cells transiently co-expressing cAMP biosensor CEpac and 5-HT7R-eGFP or 5-HT1AR-mCherry. (E, F) Representative traces showing the change in the fluorescence ratio in response to the agonist LP-211 (n≥30) and 5-HT (n≥30). Experiments were performed in cells transiently co-expressing cAMP biosensor CEpac and 5-HT7R-eGFP and 5-HT1AR-mCherry. Initial values of the fluorescence ratio were normalized to 1. Data are the means ± s.e.m of at least three separate experiments.

5-HT1AR is known to down-regulate cAMP production. To activate 5-HT1AR pre-stimulation using FSK is required to increase cAMP production. Increase in cAMP was rather transient and stabilized by adding PDE inhibitor IBMX. To identify optimal condition for activating 5-HT1AR, a set of control experiments with varying FSK and IBMX concentrations were performed (Fig. S3A). An optimal concentration for 5 µM FSK and 50 µM IBMX were chosen to obtain sufficient increase in cAMP production to activate 5-HT1AR leading to significant decrease in cAMP. The optimal concentration did not saturate the biosensor and did not suppress the 5-HT1AR effect. FSK and IBMX responses were individually controlled in the presence of CEpac only, CEpac with 5-HT7R, CEpac with 5-HT1AR, and CEpac with 5-HT7R -5-HT1AR to ensure no additional effect as shown (Fig. S3B). The ability of 5-HT1AR-mediated signaling via Gi to inhibit the FSK-induced cAMP accumulation following the receptor stimulation with LP-211 were subsequently analyzed. Negligible increase of the A/D ratio of CEpac was observed upon LP-211 stimulation, which indicated no 5-HT1AR-mediated regulation of intracellular [cAMP] (Fig. 3C). Pretreatment with the selective 5-HT1AR antagonist WAY100635 (WAY, purple trace) fully blocked the small response. Statistical evaluation of the time dependence of the A/D ratio data with the two exponential fit model no. 2 (Prasad et al., 2019) revealed a shorter decay time and smaller amplitude (*τ2*=75 ± 2.0 s, *A2*=0.08 ± 0.01, N=2, n=30) for LP-211 responses (Fig. 3D) in comparison to 5-HT7R. The kinetic properties such as amplitude and decay time help in determining the strength and speed of serotonergic signaling.

We investigated the overall cAMP signaling on superposition of both 5-HT7R and 5-HT1AR signaling cascades. Having the knowledge of the individual signaling mechanism of 5-HT7R and 5-HT1AR from earlier results, we addressed the question of whether the interaction between receptors impacts the signaling of the individual 5-HTR or if the overall signaling is a superposition of both signaling cascades. To answer that, 5-HT7R–eGFP and 5-HT1AR–mCherry together with CEpac were co-expressed in N1E cells. Upon the application of LP-211, which preferentially activates 5-HT7R, a strong variation in the CEpac signal was observed (Fig. 3E), in which an increase of [cAMP] predominated. In contrast application of 5-HT, a universal activator of serotonin receptors, a decrease in [cAMP] predominated, although the response showed a strong variability (Fig. 3F).

Statistical evaluation of the experimental data with the double exponential fit model no. 3 (Prasad et al., 2019) did not provide a conclusive result due to highly superimposing processes of cAMP up- and down-regulation.

Based on the data provided in Prasad et al., 2019 where we showed that on blocking the 5-HT1AR with WAY, the 5-HT7R-mediated response to 8-OH-DPAT provides an upregulation of [cAMP], we expected the same response in the cells co-expressing 5-HT7R and 5-HT1AR showing an upregulation of [cAMP] for the 5-HT7R-mediated response to LP-211 similar to that seen earlier in the cells that only express the 5-HT7R and CEpac. As described in Prasad et al., 2019, blocking of the 5-HT7R with SB led to a significant reduction of the expected downregulation of the FSK-induced cAMP accumulation induced by 5-HT1AR activation via Gi with 5-CT, we expected complete reduction of the expected downregulation of the FSK-induced cAMP accumulation induced by 5-HT1AR activation with LP-211 similar to that seen earlier in the cells that only express the 5-HT1AR and CEpac. To explain the diversity in cAMP response observed in (Fig. 3E) the 5-HT1AR-mediated cAMP signaling (i.e. reduction in the cAMP downregulation) must be completely suppressed because the hetero-oligomerization (5-HT7R:5-HT1AR stoichiometry) seem to be shifted more towards the 5-HT1AR, the more it forms oligomers with 5-HT7R, hence readily stimulated by LP-211. The heterogeneity in cAMP response seen in (Fig. 3F) suggests it forms oligomers more with 5-HT1AR (Renner et al., 2012) explaining the stronger cAMP downregulation.

Thus, the above data demonstrates that the ligand LP-211, which shows suitable pharmacokinetic properties, is a highly selective agonist for characterizing the functional role of 5-HT7 receptor such as the activation of the cAMP response over 5-HT1A receptor.

### Protein kinase C alpha (PKCα) activation in N1E cells

PKCα activation upon stimulation by PMA in N1E cells was characterized by performing FRET measurement at a single cell level using confocal imaging. Confocal images of cells co-expressing PKCα tagged CFP and PKCα tagged YFP plasmids showed PKCα expression in the cytosol (Fig. 4A). Cells were treated with PMA for 15 minutes which led to increase in FRET efficiency *Ef_DA50_* (6 ± 1.0 %) and fluorescence intensity at the surface in comparison to before (Fig. 4A, B). This demonstrates physical translocation of PKCα from the cytosol to the cell membrane indicating protein dimerization in the membrane. Spectral based linear unmixing (lux-FRET) measurements were performed in cell population co-expressing PKCα-CFP and PKCα-YFP at excitation range 400 nm - 520 nm at spectrofluorometer. The fluorescence intensity after PMA application changed seen through unmixing of the sample spectra at 440 nm excitation (Fig. 4C, D). The emission spectra measured at 440 nm excitation were unmixed on fitting by a linear combination of reference spectra PKCα-CFP (D) and PKCα-YFP (A) which were obtained separately. Cells showed FRET efficiency *Ef_D_* (7 ± 2.0 %) with *x_D_* value 0.5 before and after PMA application FRET reduced to (5 ± 2.0 %).

**Figure 4.**
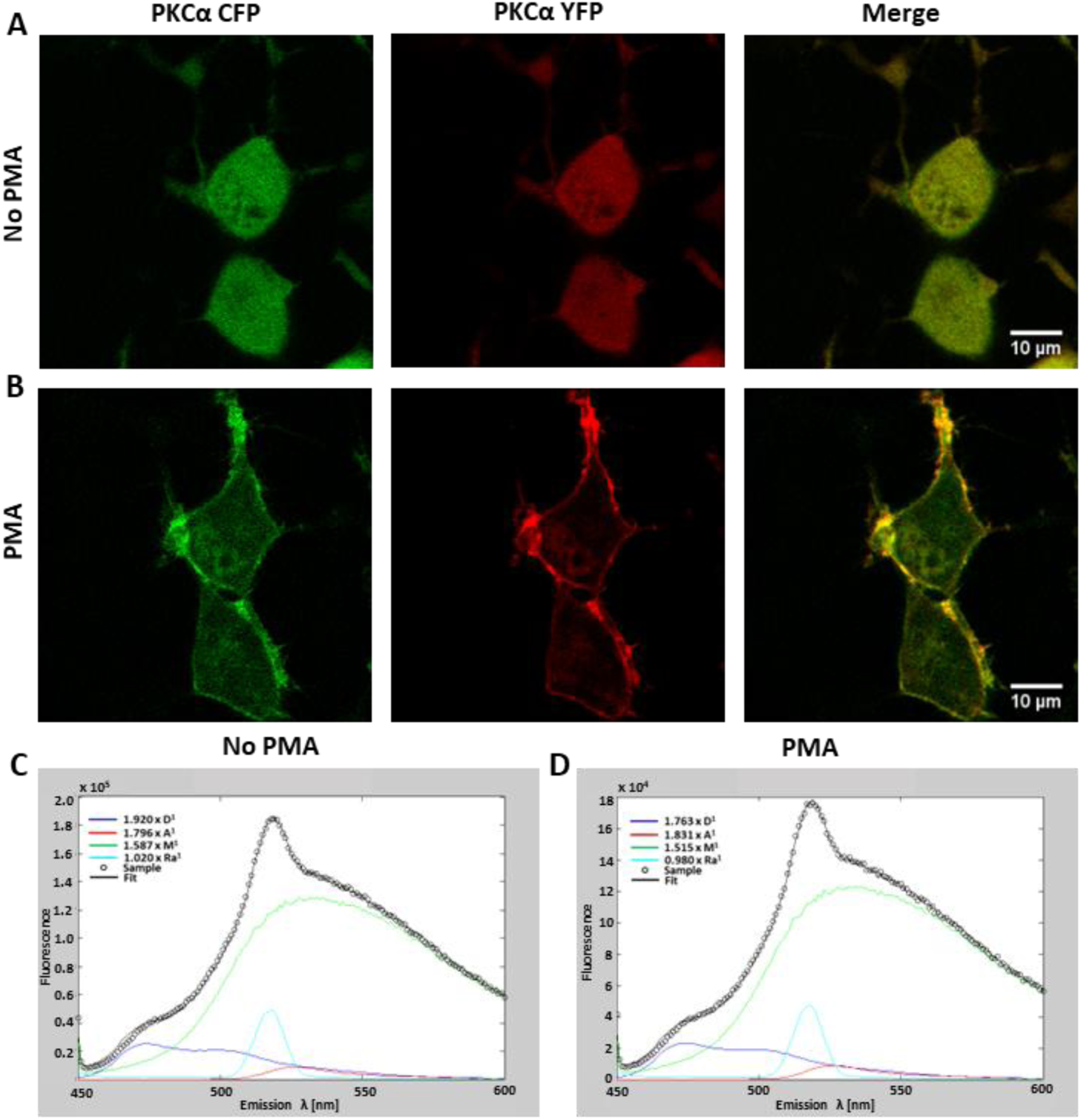
PMA induced activation of PKC-alpha in N1E cells. (A, B) Representative confocal image of a cell transiently co-expressing PKCα-CFP and PKCα-YFP. The protein was localized in the cytosol at the basal state and translocated to the plasma membrane after 15 minutes of 1 µM PMA bath application. (C, D) Unmixed sample emission spectra measured at 440 nm excitation fitted by a linear combination of reference spectra for YFP, CFP, Raman, and background before and after PMA. Experiments were repeated at least three times.

Thus, the data suggests that PKC*α* activation via PMA may activate several downward signaling pathways such as modulating 5-HT3A receptor function in the cell membrane and trafficking through an F-actin-dependent mechanism. PMA activate several PKC isoforms signal transduction pathway in many cancers, which opens the targeting of PKCs, particularly the novel PKCα, contributing to a potential avenue for therapeutic intervention.

### Physiological interaction between 5-HT receptors and CD44

CD44 acts in an antagonistic manner for the 5-HT7R. This led us to investigate the interaction between CD44 and 5-HT7 receptors and subsequently other serotonin receptors such as 5-HT1A and 5-HT4. To study that we used CD44, 5-HT7R and 5-HT1AR labeled with green and red fluorescent proteins and quantified their oligomerization by applying lux-FRET approach (Wlodarczyk et al., 2008; Zeug et al., 2012) to confocal microscopy. The spectrally highly overlapping signals were unmixed offline (see methods). Spectral based lux-FRET measurement in N1E cell population co-expressing CD44 and either 5-HT7R or 5-HT1AR or 5-HT4R were performed at spectrofluorometer.

Cells co-expressing CD44-eGFP and 5-HT receptors tagged mCherry had significantly high *Ef_D_* values (∼13% for 5-HT7R, ∼12% for 5-HT1AR) at low *x_D_* value ∼0.05 indicating only the presence of acceptor in comparison with cells co-expressing 5-HT receptors tagged eGFP and CD44-mCherry, which had significantly low *Ef_D_* (∼-0.18% for 5-HT7R, ∼-0.16% for 5-HT1AR) at high *x_D_* value ∼0.99 indicating only the presence of donor (Table S1). Cells co-expressing 5-HT1AR-eGFP and 5-HT7R-mcherry were used as a positive control. For negative control individual 5-HT receptor or tandem construct was co-expressed with eGFP or mCherry fluorescent protein showing no FRET. Positive control cells displayed apparent FRET efficiency *<u>Ef_D_</u>* ∼20% at *x_D_* 0.5 revealing strong interaction between 5-HT7R and 5-HT1AR (Table S1). Increase in fluorescence intensity in cells co-expressing CD44-eGFP with 5-HT1AR-mCherry or 5-HT7R in comparison with 5-HT4R tagged with mCherry was observed indicating a strong interaction between CD44 and 5-HT1AR and 5-HT7R (Fig. 5C-D, Table S1). The emission spectra measured at 440 nm excitation were unmixed on fitting by a linear combination of reference spectra for eGFP (D), mCherry (A). The result indicates that the N1E cells overall expressed CD44 plasmids 10 times less 5-HT1A and 5-HT7 receptors causing a strong variation in *Ef_D_* which could be due to fewer cells co-expressing CD44 plasmids with 5-HT7 or 5-HT1A receptors hence leading to a significantly low or high *x_D_* values lies depending on CD44 plasmids used as donor or acceptor. This suggests that the available donor does not couple with the available acceptor resulting in free donors or acceptors leading to artificial *Ef_D_* with low/high *x_D_*. The same measurements when performed with increased concentration transfection ratio (1:10) and (10:1) A:D for CD44 plasmids tagged with eGFP or mCherry showed improved FRET efficiency when same expression level was achieved.

**Figure 5.**
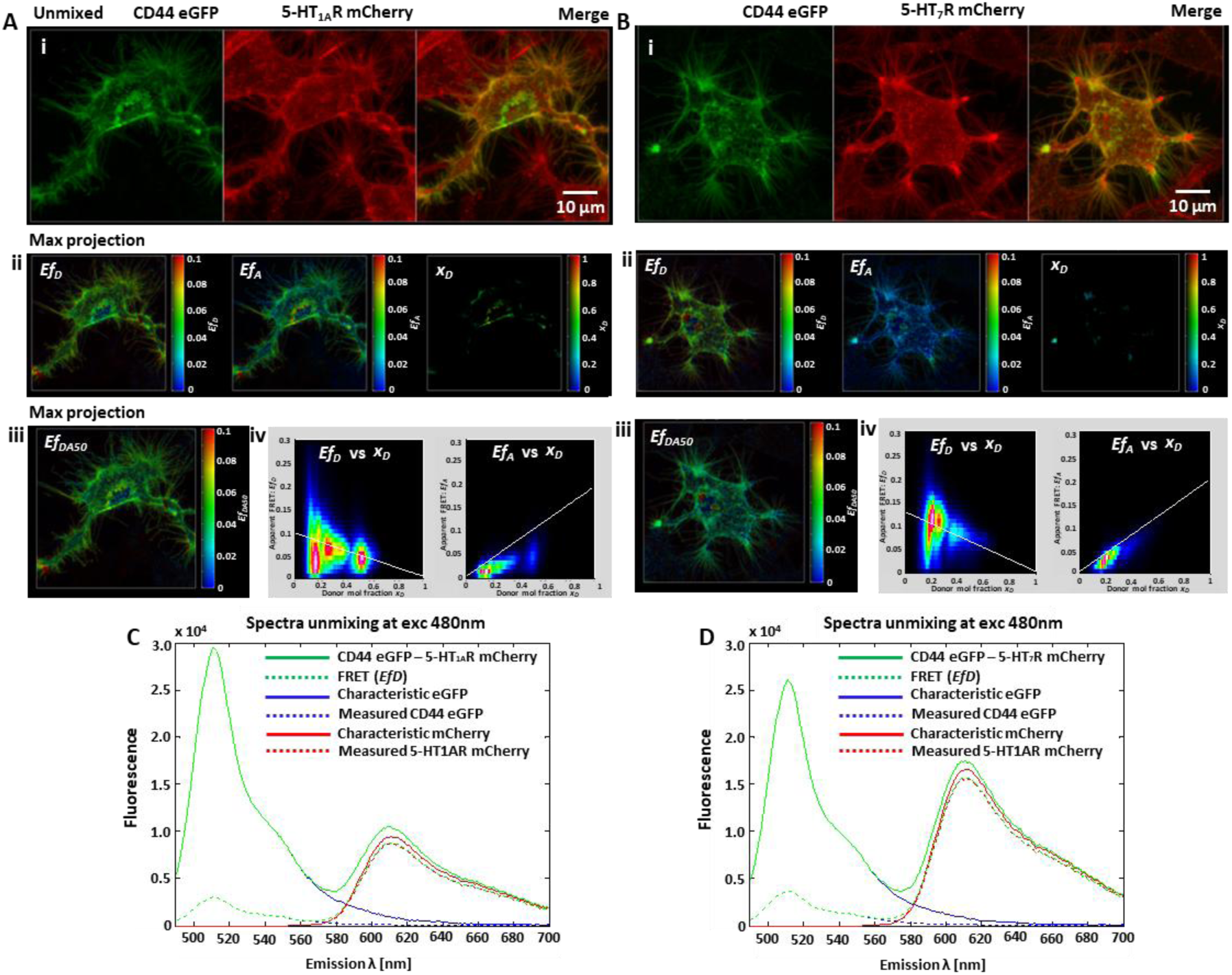
Interaction between 5-HT receptors and CD44 investigated by lux-FRET. (A, B) Representative maximum intensity z-projection of linearly unmixed images of CD44-eGFP (donor) interaction with 5-HT7 or 5-HT1A-mCherry receptors (acceptor) (i) are shown. Representative maximum intensity Z-projections are shown for apparent FRET efficiencies (*Ef_D_*, *Ef_A_*, *x_D_*) (ii) and *Ef_DA50_* (iii) of subset displaying CD44 - 5-HT receptors interaction. Representative 2D linear histogram of *Ef_D_* and *Ef_A_* (iv) are shown as functions of the donor molar fraction *x_D_* fitted by the fit model described in (Renner et al., 2012). Experimental data fitted according to the dimerization model for the FRET couples shown in A. Data were obtained from cells (n≥10). See also Figures S5 and S6. (C, D) Sample spectra unmixing fitted by a linear combination of characteristic spectra for eGFP and mCherry at excitation 480 nm showing change in the FRET efficiency.

We progressed to achieve reliable FRET efficiency for the large variation seen in cell population experiments. Lux-FRET measurements at single cell level using confocal imaging were performed. Expression profile of membrane-expressed receptors (CD44 and 5-HT) were studied. Confocal images showed that number of cells expressing 5-HT7R or 5-HT1AR in relation to CD44 was in the ratio of 10:1 (Fig. S4) confirming the earlier observation. Figure S5 shows a 2D linear histogram for selected cells co-expressing both 5-HT7R or 5-HT1AR and CD44 telling a reliable co-localization exists. Cells co-expressing 5-HT4R and CD44 showed little co-localization. Lux-FRET evaluation to confirm the interaction observed between CD44 and 5-HT receptors was performed. CD44 also localizes in Golgi explaining the staining observed in smaller region of cell surface (Fig. 5Ai, ii) (Skupien et al., 2014). Maximum intensity Z-projection of unmixed donor and acceptor in cells co-expressing 5-HT7R or 5-HT1AR tagged with mCherry and CD44 tagged with eGFP is shown (Fig. 5Ai, Bi). Maximum intensity projections of subset of cells displaying CD44 - 5-HT7 and 5-HT1A receptors interaction showed apparent FRET efficiency *Ef_D_* 7 ± 2.0 % on average (Fig. 5Aii, Bii; S6Bi, ii). Maximum intensity Z-projections of xD corrected mean FRET efficiency was found to be *Ef_DA50_* = 7 ± 1% of subset of cells (Fig. 5Aiii, Biii, S6Ci, ii). This emphasizes the accuracy of our lux-FRET approach in confocal microscopy and supports the quality of the oligomerization studies at subcellular resolution. Pixel based 2D histogram of *Ef_D_* and *Ef_A_* over *x_D_* fitted using the dimerization model (Renner et al., 2012) for the 5-HTR’s - CD44 FRET couples is shown (Fig. 5Aiv, Biv Fig. S7Ai, ii). Maximum intensity Z-projection of cells co-expressing receptors tagged with opposite fluorophores showed significantly lower *Ef_D_* 4 ± 1.0 % on average, indicating weaker interaction (Fig. S6Ai, ii; S7Ai, ii, B). Cells co-expressing 5-HT4R and CD44 showed no FRET indicating no interaction (Fig. S7B). Positive control cells co-expressing 5- HT1AR-eGFP and 5-HR7R-mCherry displayed *Ef_D_* 6 ± 2.0 % on average (Fig. S6Biii; S7Aiii, B). Cell expressing eGFP–mCherry tandem construct with a 1:1 stoichiometry had *Ef_D_* 17 ± 3% and used as positive control (Fig. S6iv, S7Aiv, B). 2D Histograms showing distribution of apparent *Ef_D_* over *x_D_* confirmed FRET efficiency of different FRET pairs and tandem construct (Fig. S8).

During each lux-FRET measurement the correlation between FRET parameters and the intensity were controlled. In case of single 5-HT7R and 5-HT1AR, no change in FRET parameters was found in the range of pixel intensity of experimental data used. FRET efficiency was not affected by molecular crowding as the FRET pair receptors were over expressed. In case of 5-HT receptors and CD44 and 5-HT7R - 5-HT1AR (positive control), receptor concentration related change of FRET parameters in contrast to single 5-HT7R and 5-HT1AR was observed. In summary, data from Figure 5 demonstrates that the interaction between CD44 and 5-HT7R and CD44 and 5-HT1AR was stronger in comparison with other serotonin receptors such as 5-HT4R.

### Dimerization of 5-HT1A receptor and mutants

We explored the ability of 5-HT1AR and mutants to dimerize resulting from specific versus random interactions by applying lux-FRET. N1E cells were transfected with various combinations of 5-HT1AR mutant plasmids in different ratios and measurements were performed at three excitations 440 nm, 458 nm and 488 nm in cell population at spectrofluorometer. Apparent FRET efficiency *Ef_D,_* as a function of donor/acceptor ratio for WT and mutated 5-HT1A receptors was determined. Specific receptor-receptor interaction was seen from the functional dependence of both *Ef_D_* and *Ef_A_* over the donor molar fraction *x_D_*. Based on the lux-FRET analysis, using the dimerization model (Gorinski et al., 2012), we estimated the number of units participating in complex, n and obtained a best fit for the value of n. Receptors exits in dimer when n is ≥ 2.0, and exits in monomeric form when n is near 1.0 (Fig .6A). The co-expression of W175A-eCFP with Y198F-eYFP decreased the n (n=1.5) saying that the number of receptors existing in monomeric form was increased compared with WT receptors (expressed alone) (Fig. 6Ai, ii). The number of receptors existing in dimeric form increased for Y198F mutant (n=3.6) and for co-expressed mutants W175A and R176K (n=3.7) compared with WT (3.0) (Fig. 6Ai, iii). The number of receptors existing in monomeric form increased for R176K mutants compared with WT (data not shown). Receptors existing in monomeric form for double mutant (R151K+R152K) increased (n=2.1) in comparison with W175A mutant (n=2.9) (Fig. 6Aiv). Table S2 shows the FRET efficiency (E) and ‘n’ values fitted individually and globally for all the possible oligomeric interaction of 5-HT1AR mutants and wild type. This model of the dimer structure has been used in the functional studies of serotonin receptor dimer formation. An iterative computational-experimental study has demonstrated that transmembrane domains TM4 or TM5 can form an interaction interface in 5-HT1AR dimers which indicates that specific amino acid interactions maintain this interface (Gorinski et al., 2012).

**Figure 6.**
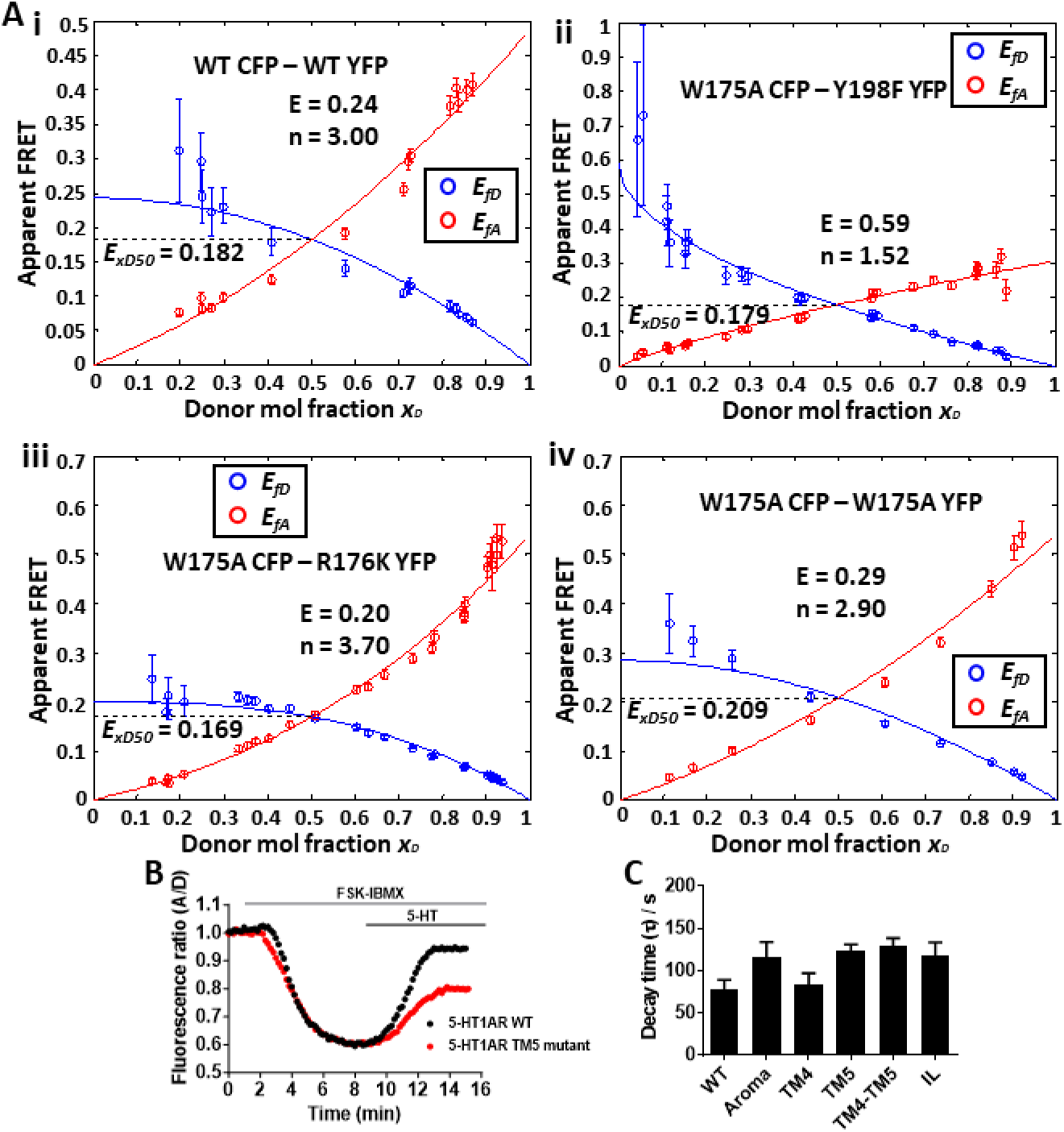
Oligomerization of 5-HT1AR WT and mutants investigated by lux-FRET. (A) Apparent FRET efficiencies *Ef_D_* (blue) and *Ef_A_* (red) were calculated and are shown as functions of the donor mole fraction *x_D_* for WT homomers (i), W175A (Aroma) - Y198F (TM5) heteromers (ii), W175A (Aroma) - R176K (TM4) heteromers (iii), W175A (Aroma) homomers (iv). Experimental data were fitted globally according to model for dynamic oligomerization described in (Renner et al., 2012; Wlodarczyk et al., 2008); where n represents a number of units participating in the complex. The total concentration of receptors was assumed to be 1. Data points represent the mean ± s.e.m from three separate experiments performed in cell populations. (B) Representative traces of the fluorescence ratio in response to agonist 5-HT in combination with FSK-IBMX stimulation (n≥20). Experiments were performed in cells transiently co-expressing cAMP biosensor CEpac and 5-HT1A WT or mutant TM5 receptor. Initial values of the ratio traces were normalized to 1. (C) The decay time of the cAMP response induced by 5-HT1AR stimulation with 5-HT, and applying fit model no2 described in (Prasad et al., 2019). Data are the means ± s.e.m of at least three separate experiments.

The importance of amino acids (Tyrosine and Trypsine) in the defined mutants of 5-HT1AR cAMP signaling pathway was further investigated using ratiometric FRET measurements. Fluorescence intensity ratio measurements were performed in cells co-expressing cAMP biosensor and either 5-HT1AR WT or mutants. In cells that expressed 5-HT1AR mutant TM5, decrease in the intracellular cAMP level on 5-HT simulation was lower than WT (Fig. 6B). Statistical evaluation of the time dependence of the fluorescence ratio data with the two exponential fit model no. 2 (Prasad et al., 2019) displayed a bigger decay time and smaller amplitude (*τ2*=123 ± 8 s, *A2*=0.20 ± 0.01, N=2, n=20) for 5-HT response in comparison with WT (*τ2*=77 ± 11 s, *A2*=0.26 ± 0.01, N=2, n=22) (Fig. 6C). Cells expressing 5-HT1AR mutant TM4 in combination with mutant TM5 showed a significantly larger decay time and smaller amplitude (*τ2*=129 ± 9 s, *A2*=0.13 ± 0.01, N=2, n=20) in comparison with WT and TM5 (Fig. 6C). Thus, the initial data shows that residues involved in 5-HT1AR mutant is important in the receptor mediated cAMP signaling.

## DISCUSSION

G protein-coupled receptors (GPCRs), including serotonin receptors, are central to the regulation of numerous cellular processes and represent key pharmacological targets in the treatment of neurological disorders. To investigate their functional interactions and signaling dynamics, we employed a combination of lux-FRET and quantitative molecular microscopy, enabling high-resolution, real-time analysis in living cells.

In this study, we applied a spectral FRET-based technique (lux-FRET) to analyze receptor interactions beyond the diffraction limit, enabling quantitative assessment of cAMP signaling, FRET efficiency, dynamic range, and specificity. Genetically encoded FRET biosensors were used to monitor intracellular cAMP levels with high precision. These biosensors are minimally affected by ion concentration changes, allowing consistent measurements across varying physiological conditions.

Cellular cAMP levels are tightly regulated by two enzyme families: adenylyl cyclases, which synthesize cAMP, and phosphodiesterases, which degrade it. Our approach provides a high-resolution, real-time platform to study these signaling dynamics and receptor interactions in living cells.

Using spectral and ratiometric FRET techniques, we assessed cAMP signaling in cells co-expressing 5-HT7R and 5-HT1AR. Selective receptor blockade with SB and WAY, followed by stimulation with partial agonists (5-HT, 5-CT, 8-OH-DPAT) revealed distinct cAMP response profiles, (Refer to Fig. 2 and Fig. S3 in Prasad et al., 2019). Treatment with a membrane-permeant cAMP precursor led to rapid intracellular metabolism and a marked shift in A/D ratio, reflecting dynamic changes in cAMP levels. Additionally, PMA-induced activation of PKC confirmed its role as a potent modulator of downstream signaling, consistent with its function as a natural activator of PKC classical isoforms.

To further dissect serotonergic signaling, we applied lux-FRET microscopy at the subcellular level to visualize receptor interactions with high spatial and temporal resolution. We observed reliable colocalization and apparent FRET efficiencies between CD44 and both 5-HT7R and 5-HT1AR, suggesting that CD44 may act as a negative regulator of 5-HT7R and functionally link extracellular matrix cues to receptor-mediated morphogenic signaling.

We also examined the oligomerization behavior of 5-HT1AR, revealing that wild-type receptors form dimers, while mutant variants showed reduced dimerization capacity. Ratiometric FRET analysis highlighted the importance of specific amino acid residues, including tryptophan and tyrosine, in stabilizing the dimeric complex. Furthermore, by analyzing homo- and hetero-oligomer formation as a function of receptor expression ratios, we demonstrated that receptor stoichiometry influences the extent of oligomerization as shown earlier in (Gorinski et al., 2012).

These findings highlight the value of studying receptor function in single living cells to complement population-level biochemical data. By combining spectral-based lux-FRET and ratiometric FRET with quantitative molecular microscopy, we investigated protein–protein interactions and receptor signaling in N1E cells. Using confocal LSM and fluorescence spectroscopy, we achieved high-resolution, real-time visualization of serotonin receptor dynamics, including oligomerization and agonist specificity, under physiological conditions and provides a powerful tool for real-time analysis of dynamic cellular processes.

## AUTHOR CONTRIBUTIONS

Performed experiments, data acquisition and collection: S.P.; Data analysis and interpretation: S.P.; Methodology and writing: S.P.

## ACKNOWLEDGMENTS

This work was supported by the Bundesministerium für Bildung und Forschung (BMBF, Federal Ministry of Education and Research, Germany, 0315690D to A.Z. and S.P.). I thank Andre Zeug for feedback.

## ABBREVIATIONS

CEpac: Cerulean/Citrine tagged exchange protein activated by cAMP
cNMP: Cyclic nucleotide mono phopshate
cAMP: Cyclic adenosine mono phosphate
cCMP: Cyclic cytidine mono phosphate
cGMP: Cyclic gunanine mono phosphate
cUMP: Cyclic uridine mono phosphate
cNMP-AM: Cyclic nucleotide mono phosphate-acetoxymethyl ester
cAMP-AM: Cyclic adenosine mono phosphate-acetoxymethyl ester
cCMP-AM: Cyclic cytidine mono phosphate-acetoxymethyl ester
cGMP-AM: Cyclic guanine mono phosphate-acetoxymethyl ester
cUMP-AM: Cyclic uridine mono phosphate-acetoxymethyl ester
FRET: Förster Resonance Energy Transfer
FSK: Forskolin
IBMX: 3-isobutyl-1-methylxanthine
Lux-FRET: linear unmixing FRET
LP-211: N-(4-cyanophenylmethyl)-4-(2-diphenyl)-1-piperazinehexanamide
PKCα: Protein kinase C alpha
PMA: 4 beta-phorbol 12-myristate 13-acetate
PO4-AM3: Phosphate tris (acetoxymethyl) ester
WT: Wild type
5-HT1AR: Serotonin receptor subtype 1A
5-HT7R: Serotonin receptor subtype 7

## REFERENCES

Beste, K. Y., & Seifert, R. (2013). cCMP, cUMP, cTMP, cIMP and cXMP as possible second messengers: development of a hypothesis based on studies with soluble guanylyl cyclase alpha(1)beta(1). Biol Chem, 394(2), 261–270. doi:10.1515/hsz-2012-0282

Bijata, M., Labus, J., Guseva, D., Stawarski, M., Butzlaff, M., Dzwonek, J., . . . Ponimaskin, E. (2017). Synaptic Remodeling Depends on Signaling between Serotonin Receptors and the Extracellular Matrix. Cell Rep, 19(9), 1767–1782. doi:10.1016/j.celrep.2017.05.023

Butzlaff, M., Weigel, A., Ponimaskin, E., & Zeug, A. (2015). eSIP: A Novel Solution-Based Sectioned Image Property Approach for Microscope Calibration. PLoS One, 10(8), e0134980. doi:10.1371/journal.pone.0134980

Davis, J. B., Calvert, V., Roberts, S., Bracero, S., Petricoin, E., & Couch, R. (2018). Induction of nerve growth factor by phorbol 12-myristate 13-acetate is dependent upon the mitogen activated protein kinase pathway. Heliyon, 4(5), e00617. doi:10.1016/j.heliyon.2018.e00617

Francis, S. H., Busch, J. L., Corbin, J. D., & Sibley, D. (2010). cGMP-dependent protein kinases and cGMP phosphodiesterases in nitric oxide and cGMP action. Pharmacol Rev, 62(3), 525–563. doi:10.1124/pr.110.002907

Gloerich, M., & Bos, J. L. (2010). Epac: defining a new mechanism for cAMP action. Annu Rev Pharmacol Toxicol, 50, 355–375. doi:10.1146/annurev.pharmtox.010909.105714

Gorinski, N., Kowalsman, N., Renner, U., Wirth, A., Reinartz, M. T., Seifert, R., . . . Niv, M. Y. (2012). Computational and experimental analysis of the transmembrane domain 4/5 dimerization interface of the serotonin 5-HT(1A) receptor. Mol Pharmacol, 82(3), 448–463. doi:10.1124/mol.112.079137

Hedlund, P. B., Leopoldo, M., Caccia, S., Sarkisyan, G., Fracasso, C., Martelli, G., . . . Perrone, R. (2010). LP-211 is a brain penetrant selective agonist for the serotonin 5-HT(7) receptor. Neurosci Lett, 481(1), 12–16. doi:10.1016/j.neulet.2010.06.036

Klarenbeek, J., & Jalink, K. (2014). Detecting cAMP with an Epac-Based FRET Sensor in Single Living Cells. In J. Zhang, Q. Ni, & R. H. Newman (Eds.), Fluorescent Protein-Based Biosensors: Methods and Protocols (pp. 49–58). Totowa, NJ: Humana Press.

Kobe, F., Renner, U., Woehler, A., Wlodarczyk, J., Papusheva, E., Bao, G., . . . Ponimaskin, E. (2008). Stimulation- and palmitoylation-dependent changes in oligomeric conformation of serotonin 5-HT1A receptors. Biochim Biophys Acta, 1783(8), 1503–1516. doi:10.1016/j.bbamcr.2008.02.021

Liu, M., Clarke, C. J., Salama, M. F., Choi, Y. J., Obeid, L. M., & Hannun, Y. A. (2017). Co-ordinated activation of classical and novel PKC isoforms is required for PMA-induced mTORC1 activation. PLoS One, 12(9), e0184818. doi:10.1371/journal.pone.0184818

Meyer, B. H., Segura, J. M., Martinez, K. L., Hovius, R., George, N., Johnsson, K., & Vogel, H. (2006). FRET imaging reveals that functional neurokinin-1 receptors are monomeric and reside in membrane microdomain. Proc Natl Acad Sci U S A, 103(7), 2138–2143.

Mogha, A., Guariglia, S. R., Debata, P. R., Wen, G. Y., & Banerjee, P. (2012). Serotonin 1A receptor-mediated signaling through ERK and PKCalpha is essential for normal synaptogenesis in neonatal mouse hippocampus. Transl Psychiatry, 2, e66. doi:10.1038/tp.2011.58

Prasad, S., Ponimaskin, E., & Zeug, A. (2019). Serotonin receptor oligomerization regulates cAMP-based signaling. J Cell Sci, 132(16). doi:10.1242/jcs.230334

Prasad, S., Zeug, A., & Ponimaskin, E. (2013). Analysis of receptor-receptor interaction by combined application of FRET and microscopy. Methods Cell Biol, 117, 243–265. doi:10.1016/B978-0-12-408143-7.00014-1

Renner, U., Zeug, A., Woehler, A., Niebert, M., Dityatev, A., Dityateva, G., . . . Ponimaskin, E. G. (2012). Heterodimerization of serotonin receptors 5-HT1A and 5-HT7 differentially regulates receptor signalling and trafficking. J Cell Sci, 125(10), 2486–2499.

Roszkowska, M., Skupien, A., Wojtowicz, T., Konopka, A., Gorlewicz, A., Kisiel, M., . . . Dzwonek, J. (2016). CD44: a novel synaptic cell adhesion molecule regulating structural and functional plasticity of dendritic spines. Mol Biol Cell, 27(25), 4055–4066. doi:10.1091/mbc.E16-06-0423

Salonikidis, P. S., Niebert, M., Ullrich, T., Bao, G., Zeug, A., & Richter, D. W. (2011). An ion-insensitive cAMP biosensor for long term quantitative ratiometric fluorescence resonance energy transfer (FRET) measurements under variable physiological conditions. J Biol Chem, 286(26), 23419–23431. doi:10.1074/jbc.M111.236869

Salonikidis, P. S., Zeug, A., Kobe, F., Ponimaskin, E., & Richter, D. W. (2008). Quantitative measurement of cAMP concentration using an exchange protein directly activated by a cAMP-based FRET-sensor. Biophys J, 95(11), 5412–5423. doi:10.1529/biophysj.107.125666

Senbanjo, L. T., & Chellaiah, M. A. (2017). CD44: A Multifunctional Cell Surface Adhesion Receptor Is a Regulator of Progression and Metastasis of Cancer Cells. Front Cell Dev Biol, 5, 18. doi:10.3389/fcell.2017.00018

Speranza, L., Labus, J., Volpicelli, F., Guseva, D., Lacivita, E., Leopoldo, M., . . . Ponimaskin, E. (2017). Serotonin 5-HT7 receptor increases the density of dendritic spines and facilitates synaptogenesis in forebrain neurons. J Neurochem, 141(5), 647–661. doi:10.1111/jnc.13962

Sun, H., Hu, X. Q., Moradel, E. M., Weight, F. F., & Zhang, L. (2003). Modulation of 5-HT3 receptor-mediated response and trafficking by activation of protein kinase C. J Biol Chem, 278(36), 34150–34157. doi:10.1074/jbc.M303584200

Taylor, S. S., Yang, J., Wu, J., Haste, N. M., Radzio-Andzelm, E., & Anand, G. (2004). PKA: a portrait of protein kinase dynamics. Biochim Biophys Acta, 1697(1-2), 259–269. doi:10.1016/j.bbapap.2003.11.029

van der Krogt, G. N., Ogink, J., Ponsioen, B., & Jalink, K. (2008). A comparison of donor-acceptor pairs for genetically encoded FRET sensors: application to the Epac cAMP sensor as an example. PLoS One, 3(4), e1916. doi:10.1371/journal.pone.0001916

Veatch, W., & Stryer, L. (1977). The dimeric nature of the gramicidin A transmembrane channel Conductance and fluorescence energy transfer studies of hybrid channels. J Mol Biol, 113(1), 89–102.

Werner, K., Schwede, F., Genieser, H. G., Geiger, J., & Butt, E. (2011). Quantification of cAMP and cGMP analogs in intact cells: pitfalls in enzyme immunoassays for cyclic nucleotides. Naunyn Schmiedebergs Arch Pharmacol, 384(2), 169–176. doi:10.1007/s00210-011-0662-6

Wlodarczyk, J., Woehler, A., Kobe, F., Ponimaskin, E., Zeug, A., & Neher, E. (2008). Analysis of FRET signals in the presence of free donors and acceptors. Biophys J, 94(3), 986–1000. doi:10.1529/biophysj.107.111773

Woehler, A., Wlodarczyk, J., & Ponimaskin, E. G. (2009). Specific oligomerization of the 5-HT1A receptor in the plasma membrane. Glycoconj J, 26(6), 749–756. doi:10.1007/s10719-008-9187-8

Wolter, S., Golombek, M., & Seifert, R. (2011). Differential activation of cAMP- and cGMP-dependent protein kinases by cyclic purine and pyrimidine nucleotides. Biochem Biophys Res Commun, 415(4), 563–566. doi:10.1016/j.bbrc.2011.10.093

