## Supplemental Material for "Quantitative spectral Linear Unmixing and Ratiometric FRET for live-cell imaging of protein interactions"

**SUPPLEMENTARY DATA**

**Table legends**

| **Fluorolog: Exc1 / Exc2 (480 nm / 580 nm)** | *Etc* | *Rtc* | |
|---|---|---|---|
| eGFP-mCherry TC | 21.14% | 30.66% | |
| **FRET couples** | $Ef_D$ | $Ef_A$ | $x_D$ |
| 5-HT$_{1A}$R eGFP - 5-HT$_7$R mCherry (+ve ctrl) | 21.26% | 16.96% | 0.44 |
| 5-HT$_7$R eGFP - 5-HT$_{1A}$R mCherry | 16.28% | 24.21% | 0.60 |
| CD44 eGFP - 5-HT$_{1A}$R mCherry | 11.71% | 0.52% | 0.04 |
| CD44 eGFP - 5-HT$_7$R mCherry | 13.23% | 0.29% | 0.02 |
| 5-HT$_{1A}$R eGFP - CD44 mCherry | -0.16% | -1.17% | 0.88 |
| 5-HT$_7$R eGFP - CD44 mCherry | -0.18% | -8.89% | 0.98 |
| CD44 eGFP - 5-HT$_4$R mCherry | 11.77% | 0.52% | 0.06 |
| 5-HT$_4$R eGFP - CD44 mCherry | 0% | 0% | 0.32 |

**Table S1. FRET efficiency shown for interaction of CD44-5-HT receptors.** Table describing interaction of CD44 with 5-HT7R or 5-HT1AR displayed through apparent FRET efficiency investigated by lux-FRET. Values are mean of at least three separate experiments.

**Individual fitting**

| **Exc1/Exc2** | **E(D)** | **E(A)** | **n(D)** | **n(A)** | **$E_{XD50}$** |
|---|---|---|---|---|---|
| **Aroma CFP - Aroma YFP** | | | | | |
| 458/488 | 0.33 | 0.23 | 2.40 | 2.50 | 0.20 |
| **IL CFP - IL YFP** | | | | | |
| 458/488 | 0.46 | 0.35 | 1.92 | 2.41 | 0.21 |
| **Aroma CFP - TM4 YFP** | | | | | |
| 458/488 | 0.21 | 0.19 | 3.42 | 3.49 | 0.16 |
| **Aroma CFP - TM5 YFP** | | | | | |
| 458/488 | 0.67 | 0.50 | 1.43 | 1.63 | 0.17 |
| **WT CFP - WT YFP** | | | | | |
| 458/488 | 0.27 | 0.22 | 2.61 | 3.25 | 0.18 |
| **TM4 CFP - TM4 YFP** | | | | | |
| 458/488 | 0.67 | 0.50 | 1.43 | 1.63 | 0.17 |
| **TM5 CFP - TM5 YFP** | | | | | |
| 458/488 | 0.21 | 0.19 | 3.42 | 3.95 | 0.16 |

**Global fitting**

| **Exc1/Exc2** | **E** | **n** | **$E_{XD50}$** |
|---|---|---|---|
| **Aroma CFP - Aroma YFP** | | | |
| 458/488 | 0.29 | 2.90 | 0.20 |
| **IL CFP - IL YFP** | | | |
| 458/488 | 0.43 | 2.07 | 0.22 |
| **Aroma CFP - TM4 YFP** | | | |
| 458/488 | 0.20 | 3.70 | 0.16 |
| **Aroma CFP - TM5 YFP** | | | |
| 458/488 | 0.59 | 1.52 | 0.17 |
| **WT CFP - WT YFP** | | | |
| 458/488 | 0.24 | 3.00 | 0.18 |
| **TM4 CFP - TM4 YFP** | | | |
| 458/488 | 0.59 | 1.52 | 0.17 |
| **TM5 CFP - TM5 YFP** | | | |
| 458/488 | 0.20 | 3.70 | 0.16 |

**Table S2. Oligomerization of various mutants of 5-HT1A receptor.** Table describing oligomerization between FRET couples 5-HT1AR WT and mutants investigated by spectral lux-FRET displaying apparent FRET efficiency fitted individually and globally. Values are mean of at least three separate experiments.

### Figure legends

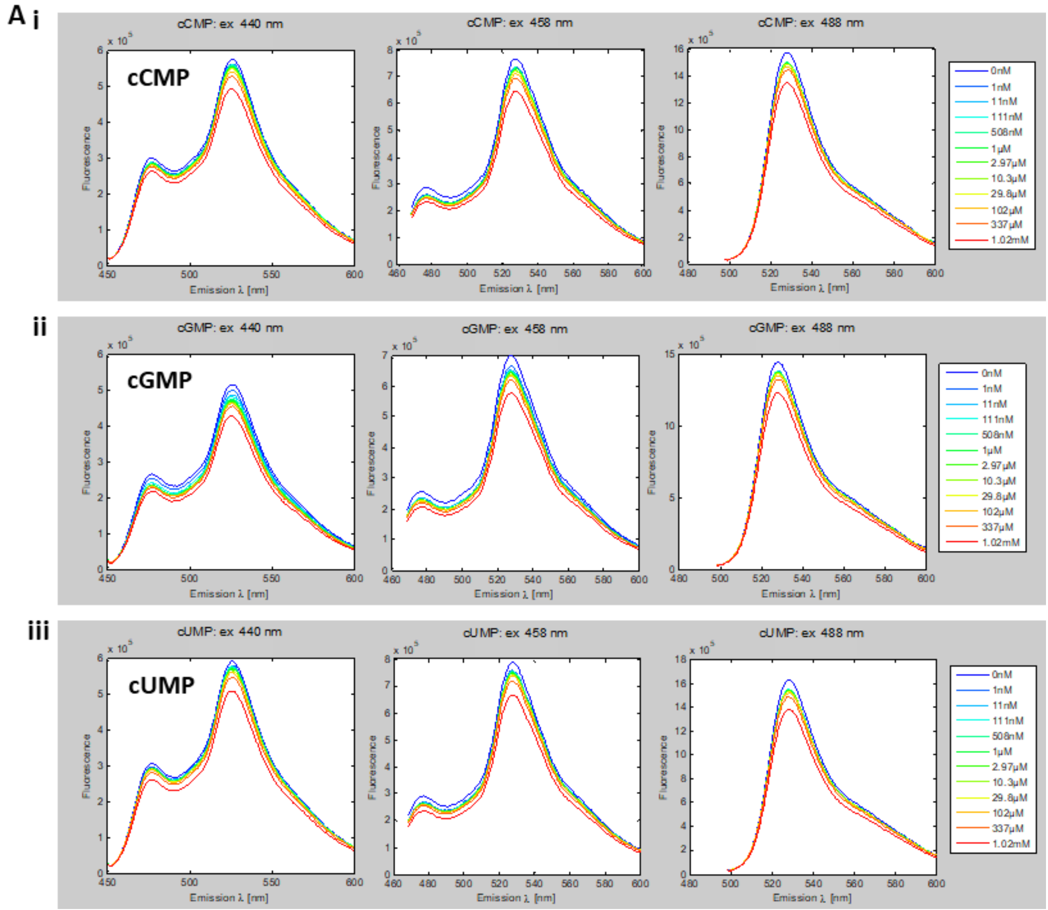


**Figure S1. Cyclic nucleotides titration to cAMP biosensor.** (A) Epac emission spectrum for cNMP, cGMP, cUMP titrations (i, ii, iii) showing no changes in the fluorescence emission signal obtained at 440 nm, 458 nm, 488 nm excitations. cNMP's were directly applied into the supernatant solution. Experiments were performed in lysed cells transiently expressing cAMP biosensor CEpac and repeated at least three times.

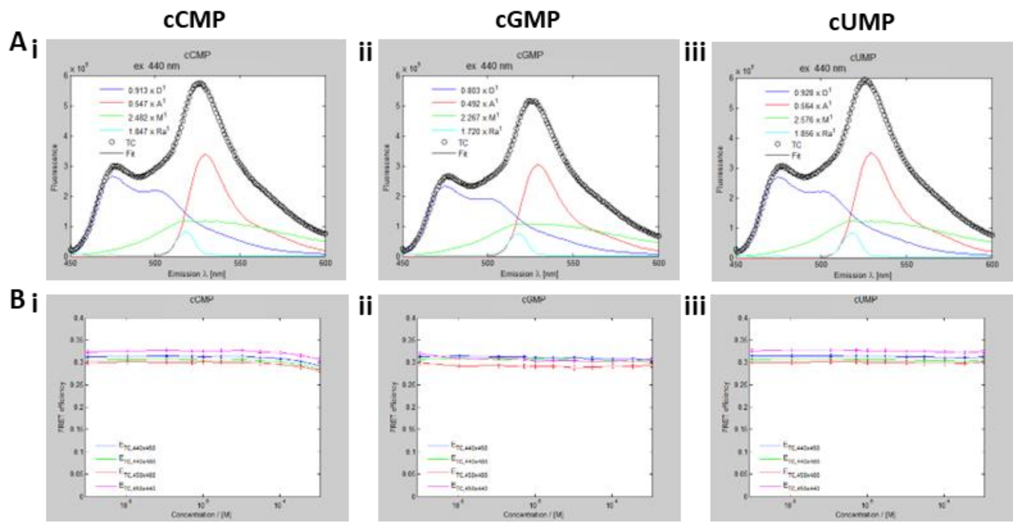


**Figure S2. Cyclic nucleotides titration to cAMP biosensor investigated by lux-FRET.** (A) Epac emission spectrum for cCMP, cGMP and cUMP measured at 440 nm excitation unmixed by a linear combination of reference spectra for YFP, CFP, Raman, and background. (B) FRET efficiency of cCMP, cGMP and cUMP at four different combinations of the three excitations over a concentration range of 0-1 mM. Experiments were performed in lysed cells transiently expressing cAMP biosensor CEpac and repeated at least three times.

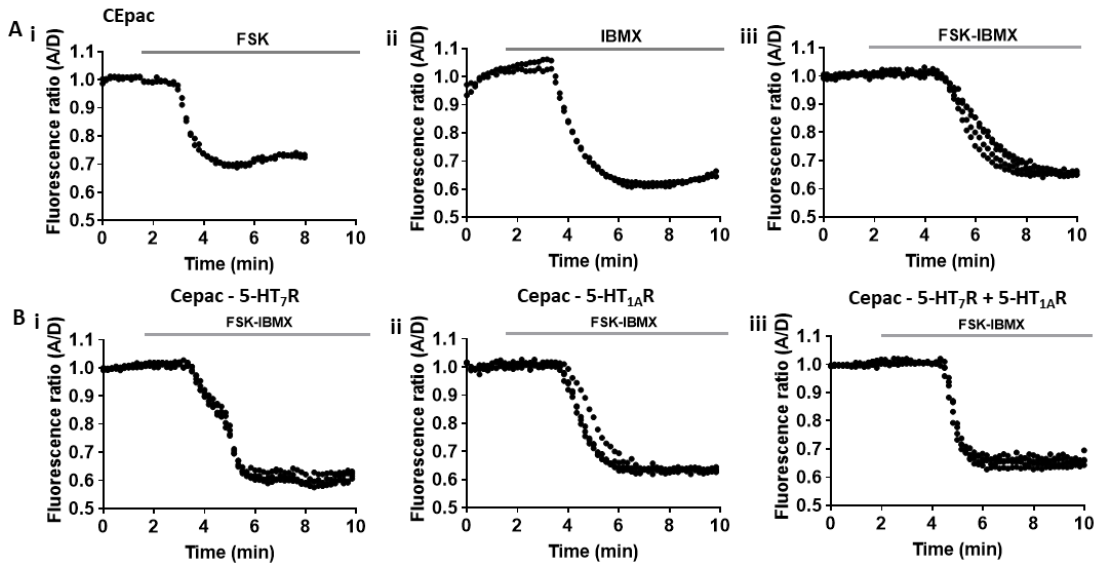


**Figure S3. FSK and IBMX induced cAMP response correspond to biosensor and receptor in N1E cells.** (A) Representative traces of the fluorescence ratio in response to 5 µM FSK and 50 µM IBMX in cells expressing biosensor. (B) Representative traces of the fluorescence ratio in response to 5 µM FSK and 50 µM IBMX in cells co-expressing biosensor with either 5-HT7R (i) or 5-HT1AR (ii) or both receptors 5-HT7R - 5-HT1AR. Initial values of the fluorescence ratio were normalized to 1.

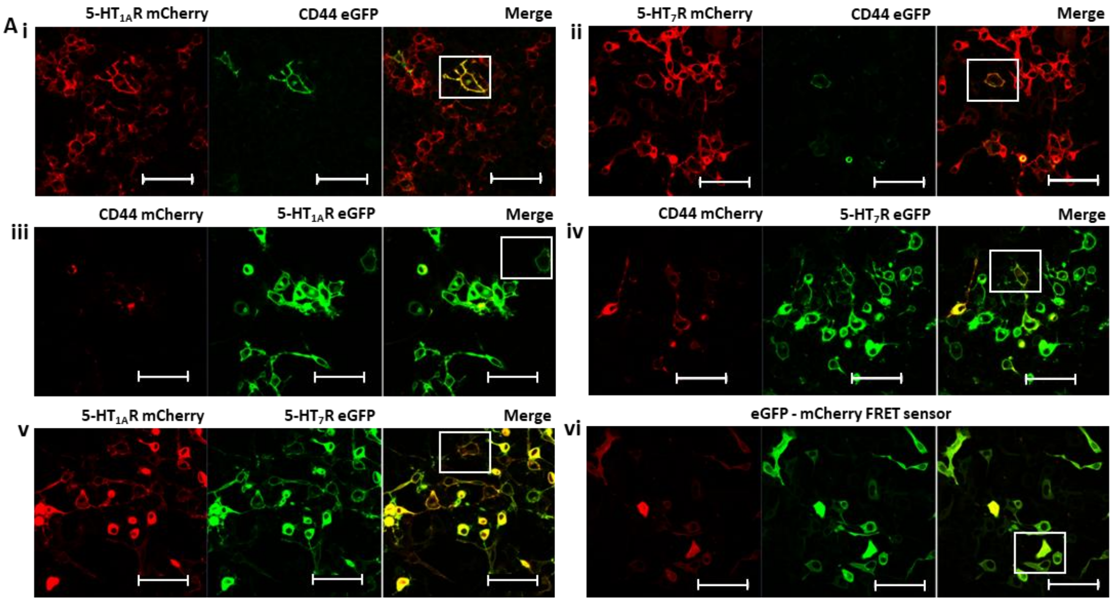


**Figure S4. Expression profile of CD44 with 5-HT receptors in N1E cells.** (A) Confocal visualization of transiently co-expressed CD44 with either 5-HT1AR (i, iii) or 5-HT7R (ii, iv) tagged with either mCherry or eGFP, 5-HT1AR and 5-HT7R co-expressed (v) which exhibits predominant localization to the plasma membrane at a transfection level of 1:1 ratio. FRET sensor eGFP - mcherry exhibiting localization to the cell body (vi). Inset showing a higher magnification view of colocalized CD44 with 5-HT receptors. Scale bar 100 µm.

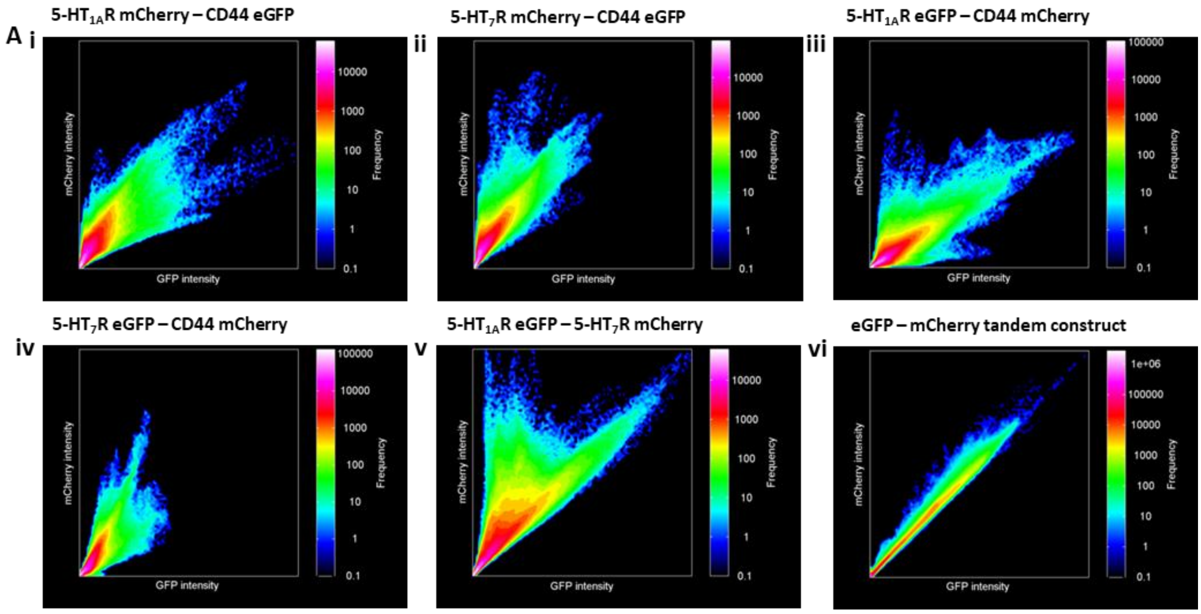


**Figure S5. Expression profile of CD44 with 5-HT receptors in N1E cells.** (A) Linear correlation fluorescence intensity histogram of donor vs acceptor for subset of cells co-expressed with CD44 receptors and 5-HT receptors tagged with mCherry or eGFP (i-iv) and 5-HT1A and 5-HT7 receptors (v) and respective tandem construct (vi) at 1:1 ratio is shown.

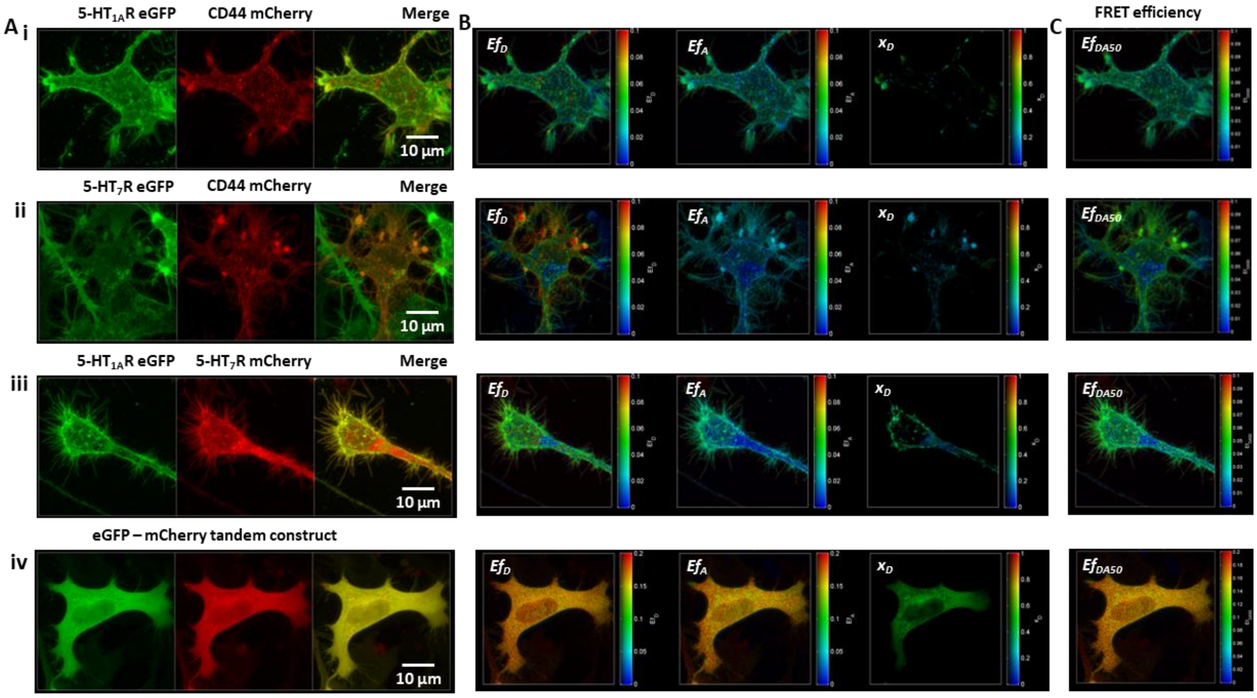


**Figure S6. Interaction between 5-HT receptors and CD44 investigated by lux-FRET.** (A) Representative maximum intensity Z-projection of linearly unmixed images of 5-HT7R or 5-HT1AR-eGFP (donor) with CD44-mCherry (acceptor) (i, ii) interaction are shown. Interactions between 5-HT7R-eGFP and 5-HT1AR-mCherry (iii) and eGFP – mCherry tandem construct (iv) shown as positive controls. (B, C) Representative maximum intensity projections of the apparent FRET efficiency ($Ef_D$, $Ef_A$, $x_D$) and $Ef_{DA50}$ of subset displaying interactions in A is shown.

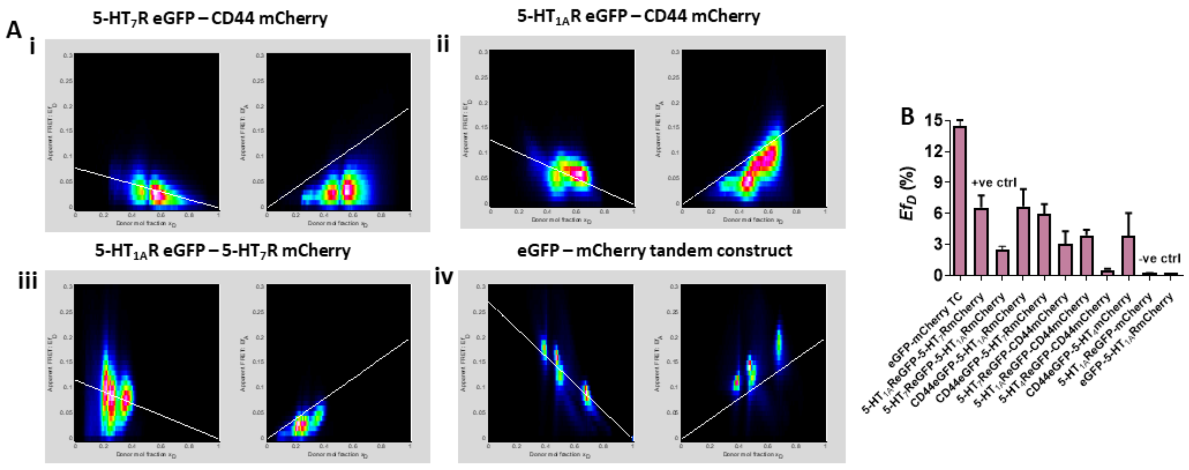


**Figure S7. Lux-FRET 2D histogram for 5-HT receptors - CD44 interaction.** (A) Representative 2D histograms of $Ef_D$ and $EfA$ over $x_D$. Experimental data fitted according to the dimerization model for the FRET couples shown in Fig. S4. (B) Quantification of FRET efficiencies ($Ef_D$) for interaction between various FRET couples CD44 and 5-HT receptors. Data were obtained from cells (n≥10).

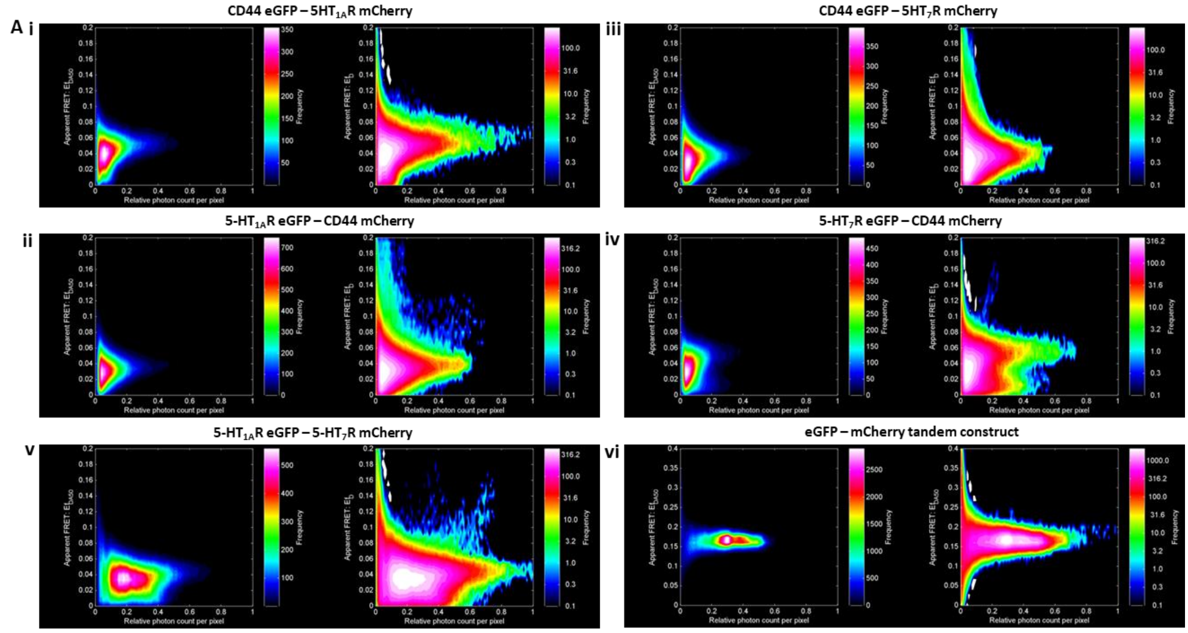


**Figure S8. 2D histogram displaying FRET efficiency based on frequency distribution.** (A) FRET quantities over relative photon count per pixel of subset based on expression level are shown for 5-HT receptors - CD44 interaction and co-expressed 5-HT receptors and related tandem construct. $Ef_{DA50}$ was calculated by having 1:1 D:A ratio at $x_D$ 0.5. Data shows linear and logarithmic sale.

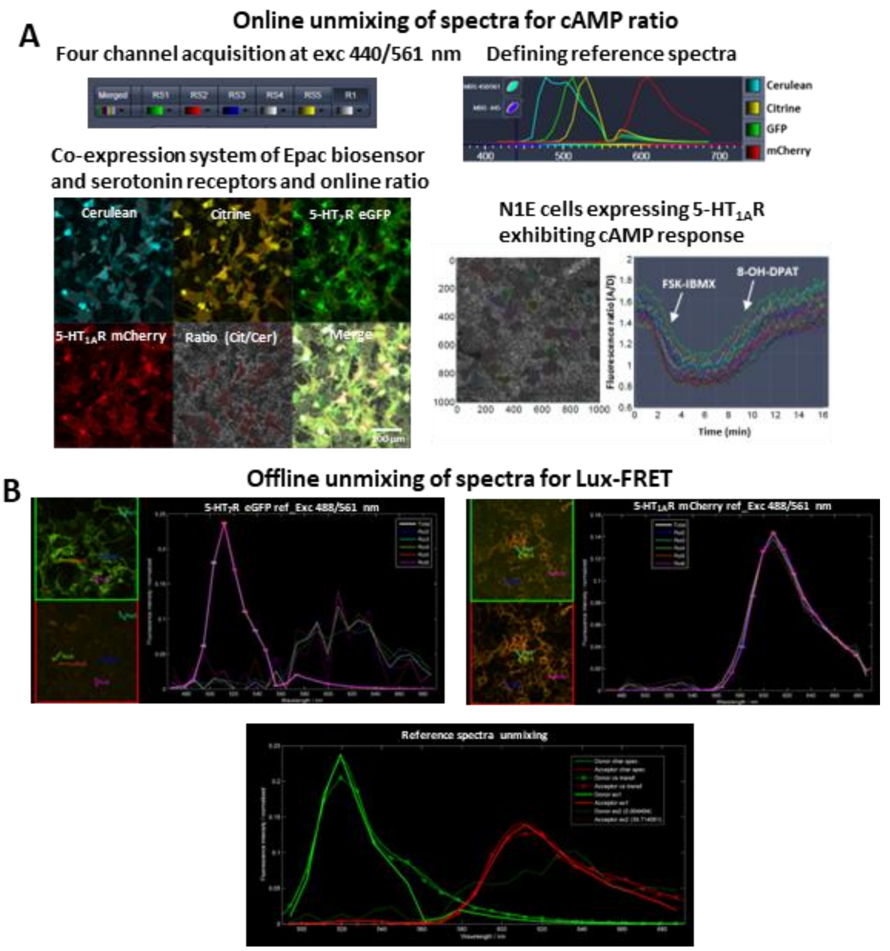


**Figure S9. Linear unmixing of reference spectra.** (A) Online unmixing of reference spectra of CEpac biosensor and 5-HT7 - eGFP and 5-HT1A - mCherry receptors expressed in N1E cells at two excitation tracks with four emission channels demonstrating change in fluorescence ratio (A/D) upon stimulation exhibiting cAMP response. (B) Offline unmixing of reference spectra of 5-HT7 - eGFP (donor) and 5-HT1A - mCherry (acceptor) receptors expressed in N1E cells at exc1 and exc2, emission range 450 – 700 nm for lux-FRET analysis.
